# Highly Dynamic and Sensitive NEMOer Calcium Indicators for Imaging ER Calcium Signals in Excitable Cells

**DOI:** 10.1101/2024.03.08.583332

**Authors:** Wenjia Gu, Jia-Hui Chen, Yiyin Zhang, Zhirong Wang, Jia Li, Sijia Wang, Hanhan Zhang, Amin Jiang, Ziyi Zhong, Jiaxuan Zhang, Chao Xi, Tingting Hou, Donald L. Gill, Dong Li, Yu Mu, Shi-Qiang Wang, Ai-Hui Tang, Youjun Wang

**Author notes:** These authors contributed equally to this work (Wenjia Gu, Jia-Hui Chen, Yiyin Zhang, Zhirong Wang, Jia Li).

## Abstract

Endoplasmic/sarcoplasmic reticulum (ER/SR) sits at the heart of the calcium (Ca^2+^) signaling machinery, yet current genetically encoded Ca^2+^ indicators (GECIs) lack the ability to detect elementary Ca^2+^ release events from ER/SR, particularly in muscle cells. Here we report a set of organellar GECIs, termed NEMOer, to efficiently capture ER Ca^2+^ dynamics with increased sensitivity and responsiveness. Compared to G-CEPIA1er, NEMOer indicators exhibit dynamic ranges that are an order of magnitude larger, which enables up to 5-fold more sensitive detection of Ca^2+^ oscillation in both non-excitable and excitable cells. The ratiometric version further allows super-resolution monitoring of local ER Ca^2+^ homeostasis and dynamics. Notably, the NEMOer-f variant enabled the inaugural detection of Ca^2+^ blinks, elementary Ca^2+^ releasing signals from the SR of cardiomyocytes, as well as *in vivo* spontaneous SR Ca^2+^ releases in zebrafish. In summary, the highly dynamic NEMOer sensors expand the repertoire of organellar Ca^2+^ sensors that allow real-time monitoring of intricate Ca^2+^ dynamics and homeostasis in live cells with high spatiotemporal resolution.

---

Calcium (Ca^2+^) signaling is indispensable for the orchestration of multiple cellular functions and physiological processes^1^. As one of the major Ca^2+^ sources and internal stores in animal cells, endoplasmic/sarcoplasmic reticulum (ER/SR) takes the center stage in the choreography of calcium signaling ^2^. Dysregulated ER homeostasis, recognized as an instigator of ER stress and apoptosis ^3^, is associated with various diseases, including cancer, cardiovascular disorders, and neurodegenerative diseases ^4,5^. To better unveil the roles of ER Ca^2+^ in health and disease, it is imperative to visualize the spatiotemporal dynamics of ER/SR Ca^2+^ with high precision and resolution.

By introducing Ca^2+^-binding site into fluorescence proteins, or via mutagenesis of Ca^2+^-binding-domain of cytosolic GECIs like Cameleon ^6^, GCaMP ^7,8^, GECO ^9^, or aequorin ^10^, researchers have developed over a dozen sensors with relatively low Ca^2+^ affinities (K_d_ > 400 μM). Notable examples of ER/SR Ca^2+^ include YC4er, CatchER sets ^11–13^, GCaMPER ^14^, CEPIA1er based series ^15–18^, and erGAP2/3 ^19^. However, these ER-resident green GECIs among them often exhibit limited dynamic ranges (<5) ^13,15^ or relatively slow kinetics^14^, thereby impeding real-time monitoring of rapid Ca^2+^-modulated events. To accurately capture elementary local ER Ca^2+^ changes^20,21^, such as Ca^2+^ blinks in cardiac muscle cells^22^, an ideal ER GECI should satisfy the following criteria: low Ca^2+^ affinity, large dynamics, and rapid kinetics.

Mostly by mutating Ca^2+^-binding residues in our recently developed NEMO indicators ^23^, we present herein a set of highly dynamic and sensitive green GECIs tailored for ER/SR, termed NEMOer, along with its ratiometric version TuNer. Basal fluorescence and responses of NEMOer indicators can be one order of magnitude larger than G-CEPIA1er, reporting significantly larger Ca^2+^ oscillation signals. TuNer sensors enable super-resolution recording of SR/ER Ca^2+^ homeostasis and dynamics. With the NEMOer-f variant, we successfully detected Ca^2+^ blink, the elementary Ca^2+^ releasing signal from SR of cardiomyocytes, as well as *in vivo* spontaneous SR Ca^2+^ releasing event in zebrafish. Collectively, the highly dynamic NEMOer sensors hold high potential for a wide range of applications in monitoring ER Ca^2+^ dynamics and homeostasis within live cells.

## Engineering of low Ca^2+^ affinity NEMOer indicators

By introducing known Ca^2+^-affinity-reducing mutations into the calmodulin domain of NEMO indicators ^15,23^ and adding ER-targeting sequences, we generated NEMO variants for monitoring ER Ca^2+^, and screened their performance in HEK293 cells (**Fig. 1A–C**, and **Supplementary Tables 1-2**). Basic sensor properties including their basal fluorescence (F_0_), maximal (F_max_) and minimal (F_min_) fluorescence were obtained in cells transiently expressing these variants at comparable levels. Two well-known ER Ca^2+^ sensors G-CEPIA1er ^15^ and ER-GCaMP6-150 ^24^ were used as controls. Briefly, after recording of F_0_, F_min_ was measured by depleting ER Ca^2+^ store with 2.5 μM ionomycin (iono; a Ca^2+^ ionophore) and subsequent permeabilization using 25 μM digitonin. In the end, large amount of Ca^2+^ (30 mM) was added to the bath containing 25 μM digitonin to induce F_max_ in permeabilized cells (**Fig. 1B**). Fluorescence responses relative to F_min_ (**Fig. 1C**), mean F_0_ and dynamic range (DR) values defined as ΔF / F_min_, or (F_max_ - F_min_) / F_min_ (**Fig. 1D**) were plotted to visualize the full capacity of these low Ca^2+^ affinity NEMOer variants.

**Figure 1.**
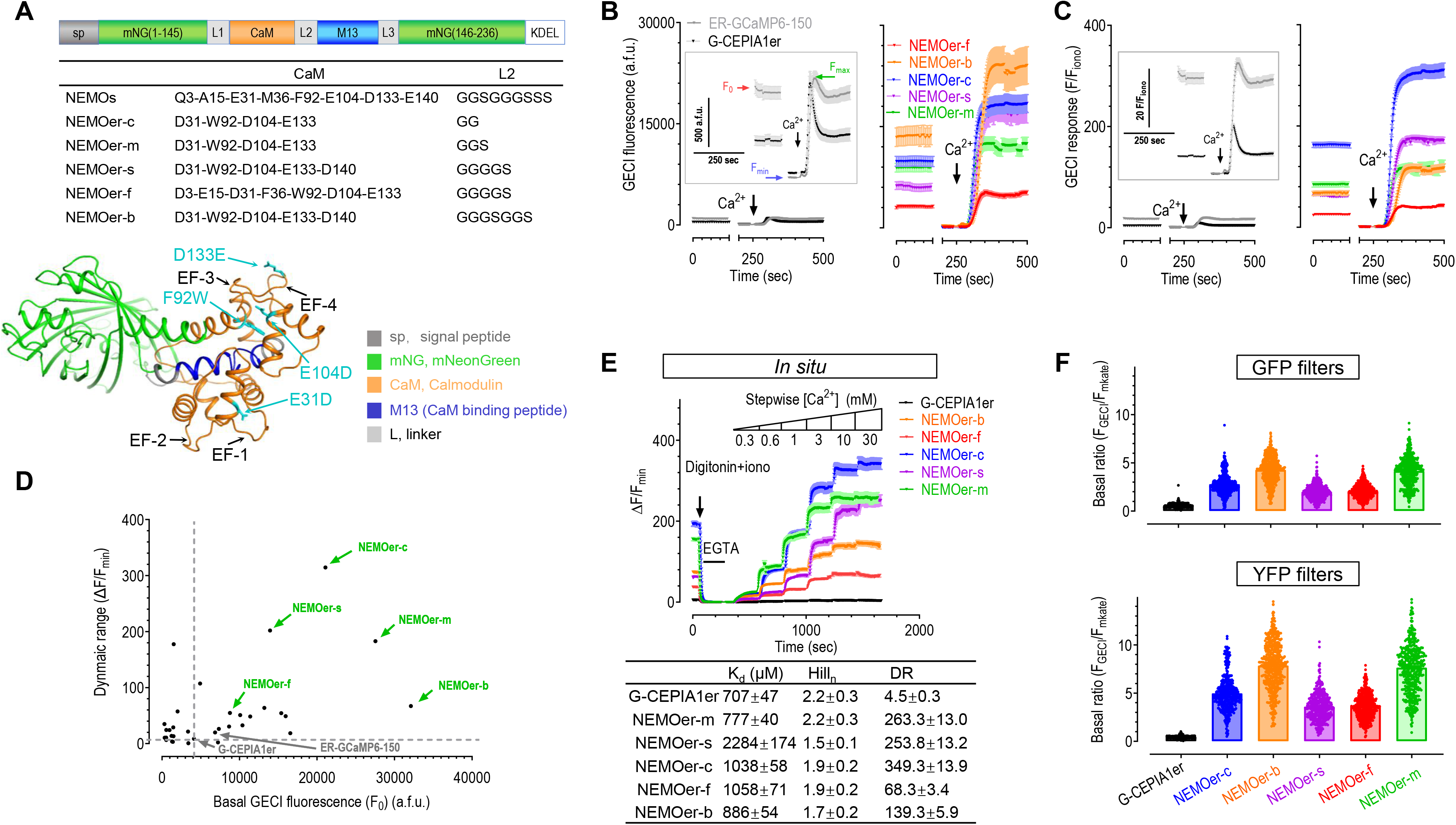
Screening and *ex vivo* characterization of NEMOer indicators. (A) NEMOer sensors are generated by introducing amino acid substitutions in calmodulin region of NEMO indicators. Top panel, a diagram showing the design of NEMOer variants. Key amino acids substitutions introduced into NEMO indicators to generate NEMOer variants are shown in a table (middle panel) or the predicted NEMOer-s structure (bottom panel) by I-TASSER ^34^ (Iterative Threading ASSEmbly Refinement). Four loops of EF-hands (EF) within calmodulin domain of NEMOer sensors are indicated by black arrows. (B–E) Screening of NEMOer variants in non-excitable mammalian cells. (B) Ca^2+^-imaging-based screening assay shown by typical traces from ER-GCaMP6-150-, G-CEPIA1er- (left) or NEMOer variants (right)-expressing cells. Inserts, traces with enlarged scales. After recording the basal fluorescence (F_0_), endoplasmic reticulum Ca^2+^ store was depleted with 2.5 μM ionomycin (iono) to obtain minimal GECI fluorescence (F_min_). Afterwards, the cells were permeablized with 30 mM Ca^2+^ imaging solution containing 25 μM digitonin to read the maximal response (F_max_). (C) Representative traces of ER-GCaMP6-150, G-CEPIA1er (left), or selected NEMOer indicators (right). (D) Scatter plot of F_0_–mean dynamic range (DR; (F_max_ - F_min_) / F_min_) of the indicated GECIs. (E) *In situ* dose–response curves of NEMOer sensors. Top, typical traces; Bottom, statistics (three independent biological replicates; >17 cells per repeat). Data shown as mean ± s.e.m. (F) Basal brightness of NEMOer or G-CEPIA1er sensors viewed with YFP (top) or GFP (bottom) filters. To achieve more accurate estimation of the basal fluorescence of GECIs (F_GECI_), F_GECI_ of cells expressing mKate-P2A-GECI constructs were normalized against the fluorescence of mKate, an expression marker (F_mKate_). (G-CEPIA1er, n = 390 cells; NEMOer- m, n = 431 cells; NEMOer-s, n = 421 cells NEMOer-f, n = 469 cells; NEMOer-c, n = 411 cells; NEMOer-b, n = 473 cell.) Three independent biological repeats. Data shown as mean ± s.e.m.

Five best-performing constructs with bright F_0_ and large DR values (**Fig. 1D**) were selected and named as m<u>Ne</u>onGreen-based Calciu<u>m</u> indicat<u>o</u>r for <u>ER</u> (NEMOer), including the medium (NEMOer-m), high contrast (NEMOer-c), fast (NEMOer-f), bright (NEMOer-b) and sensitive (NEMOer-s) versions.

We next determined performance of NEMOer variants both *in vitro* and *in situ.* The affinities of NEMOer variants were reduced to near milimolar (mM) ranges (**Extended Data Fig. 1A; Fig. 1E**), either comparable to that of G-CEPIA1er (707 ± 47 μM *in situ*) or significantly lower. And the Ca^2+^ dissociation kinetics of NEMOer-f (*k_off_* = 33.1 ± 0.8) is comparable to that of G-CEPIA1er (30.6 ± 8.9, **Extended Data Fig. 1B**), indicating that it may be suitable for decoding fast acting ER Ca^2+^ signals in excitable cells.

More careful *in situ* characterization revealed that the *in cellulo* DR of NEMOer sensors are superior to other ER GECIs tested side-by-side. In HEK293 cells, ER-GCaMP6-150 showed a F_0_ that was close to its F_max_, indicating that it is close to saturation at the basal condition (**Fig. 1B&C**). We thus only used G-CEPIA1er as a reference as it showed a good combination of affinity, kinetics and DR^15^ (**Fig. 1B&C**). In HeLa cells, the DR values of NEMOer-f and NEMOer-b, 68.3 and 139.3, respectively, were 14.3 to 29.9-fold higher than that of G-CEPIA1er (4.5). The DRs of NEMOer-m, NEMOer-s and NEMOer-c were further increased (263.3, 253.8, or 349.3, respectively) to be over 50-fold higher than that of G-CEPIA1er (**Fig. 1E**). Overall, NEMOer indicators represent a class of bright ER Ca^2+^ indicators with the *in cellulo* DRs up to 80-fold larger than G-CEPIA1er.

All NEMOer variants showed *in vitro* DR values larger than 120-fold (**Extended Data Fig. 1A, left**), at least 18.8 fold larger than that of G-CEPIA1er (6.4 ±0.1). Similar to their corresponding template ^23^, the cause of these significantly large DR values is greater Ca^2+^-dependent fold-of-increase in the molecular brightness of the anionic fluorophores in NEMOer variants (**Extended Data Fig. 1C-G; Supplementary Table 3**). In the absence of Ca^2+^, the anionic fluorophore of NEMOer-c (0.16 ± 0.03 mM^−1^cm^−1^) is considerably dimmer, approximately one-fourth that of G-CEPIA1er. And the brightness of Ca^2+^-saturated anionic NEMOer-c (54.85 ± 2.19 mM^−1^cm^−1^), more than ten times that of G-CEPIA1er.

Consistent with the observation that the basal fluorescence (F_0_) of NEMOer sensors were much brighter than that of G-CEPIA1er (**Fig. 1D**), over 87% of G-CEPIA1er fluorophores exist in neutral state, the relative dim and less Ca^2+^-sensitive configuration, while up to 75 % of the fluorophore within NEMOer-c could exist in its bright anionic form (**Supplementary Table 3**). We next more carefully compared F_0_ of NEMOer indicators with G-CEPIA1er using a P2A-based bicistronic vector to drive the co-expression of GECIs and mKate as an expression marker at a near 1:1 ratio. The normalized basal GECI brightness was shown as the fluorescence ratio of GECI over mKate (**Fig. 1F**). As expected, all NEMOer sensors showed significantly higher basal brightness than that of G-CEPIA1er, with NEMOer-f the dimmest and NEMOer-b the brightest among all NEMOer variants. Even examined with setting optimized for G-CEPIA1er (GFP filters), F_0_ of NEMOer-f and NEMOer-b was about 3 or 8.5 fold of G-CEPIA1er, respectively. When a filter set (YFP) optimized for NEMOer was used, NEMOer-f and NEMOer-b were 7.4 and 17.4 folds brighter than G-CEPIA1er.

NEMOer indicators also showed significantly enhanced photostability than G-CEPIA1er (**Extended Data Fig. 2A&B**). It endured over 50 times (0.57 mW) higher illumination than G-CEPIA1er (0.01 mW), and showed no apparent photobleaching. The stronger illumination (from 0.01 mW to 0.57 mW) could potentially enhance the basal fluorescence of NEMOer-f sensor by over 10-fold (**Extended Data Fig. 2**), greatly broadening the applicability of NEMO sensors in scenarios requiring stronger light illumination, such as monitoring Ca^2+^ signals with super-resolution imaging system, or *in vivo* imaging of subcellular compartments such as dendrites.

Together, these results firmly establish NEMOer indicators as a class of highly bright, photostable GECIs with extraordinarily large Ca^2+^-dependent changes in fluorescence.

## Performance of NEMOer indicators in non-excitable cells

We first examined the ability of NEMOer indicators to report the depletion of ER Ca^2+^ stores induced by various stimuli. Consistent with their large DR values, in responses to full ER store depletion triggered by ionomycin, NEMOer indicators showed more than 95.8 % decreases in fluorescence, significantly larger than those form G-CEPIA1er (71.9 ± 1.3 %) (**Fig. 2A**). In response to sub-maximal activation of muscarinic acetylcholinergic receptors with carbachol (CCh, 10 μM), NEMOer indicators showed stronger response with the mean peak amplitudes 2.7 ∼ 4.1 folds higher than G-CEPIA1er (**Fig. 2B**). Similarly, NEMOer sensors also showed superior performance in reporting Ca^2+^ oscillations over G-CEPIA1er (**Fig. 2C**). To examine whether the fast NEMOer-f sensors could reliably follow cytosolic Ca^2+^ oscillations, we linked a cytosolic red GECI, R-GECO1.2 ^25^, with NEMOer-f and expressed the corresponding construct in HEK293 cells. 10 μM CCh induced synchronized changes in R-GECO1.2 and NEMOer-f fluorescence (**Fig. 2D**). Enlarged view of individual GECI transients revealed that the CCh-induced decreases in NEMOer-f signal preceded the increase in R-GECO1.2 fluorescence. This is consistent with the notion that Ca^2+^ release from ER is the Ca^2+^ source for cytosolic Ca^2+^ oscillation. Together, intensiometric NEMOer response to these investigated Ca^2+^ signals in non-excitable cells are remarkably larger than those of G-CEPIA1er tested side-by-side.

**Figure 2.**
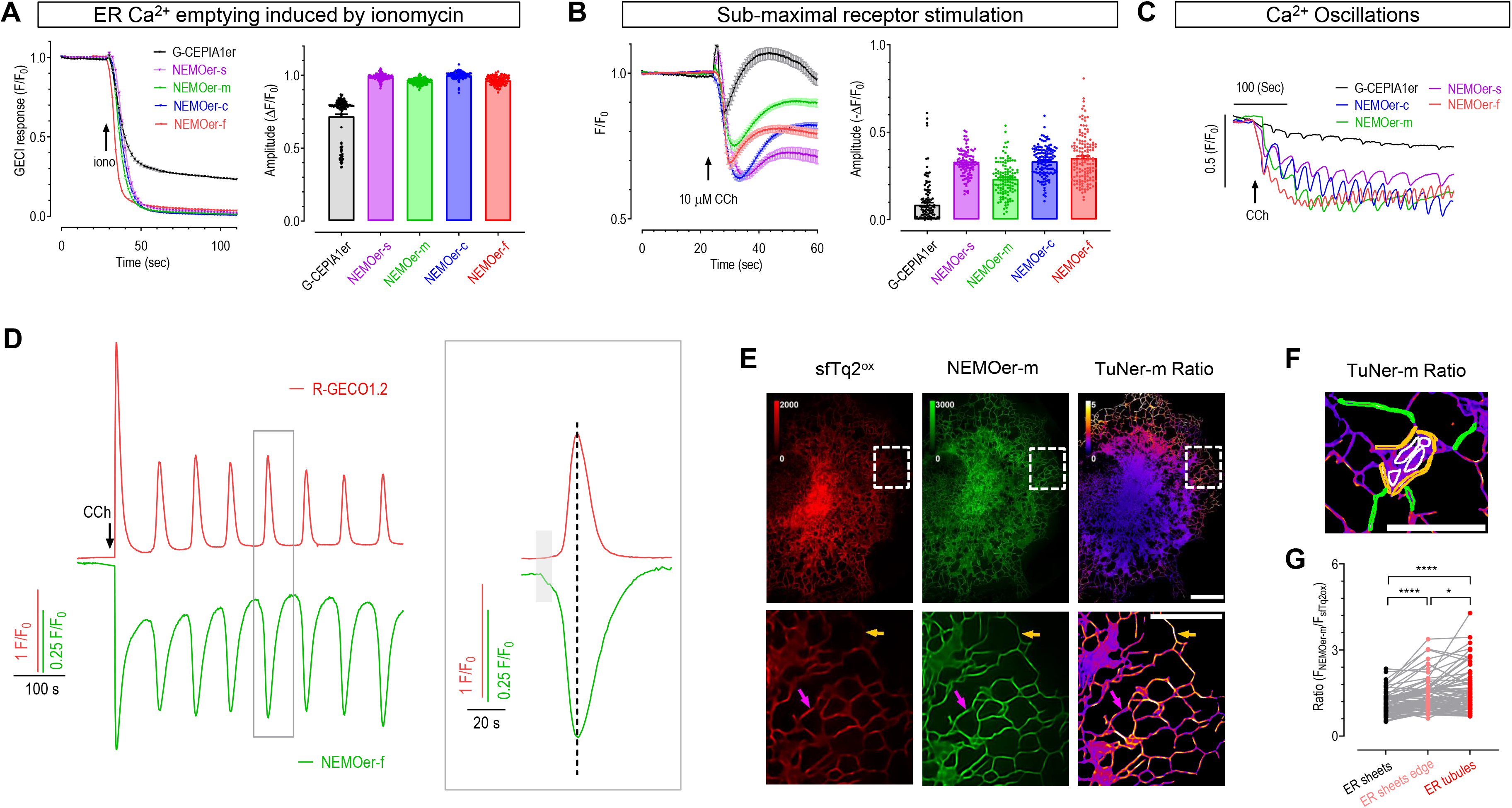
Performance of NEMOer sensors in non-excitable mammalian cells. (A) Typical ER-Ca^2+^-emptying responses in HEK293 cells induced by ionomycin (iono, 2.5 μM), as indicated by G-CEPIA1er and NEMOer indicators. (Statistics, G-CEPIA1er, n = 129 cells; NEMOer-m, n = 125 cells; NEMOer-s, n = 94 cells NEMOer-f, n = 147 cells; NEMOer-c, n = 102 cells.) n = 3 independent biological replicates, with at least 15 cells per repeat. (B-C) G-CEPIA1er or NEMOer responses to ER-Ca^2+^-lowering by submaximal receptor stimulation with carbachol (CCh, 10 μM) in HEK293 cells. Initial ER Ca^2+^ decreasing recorded in nominally Ca^2+^ free solution. Left, typical traces; right, statistics (B) (G-CEPIA1er, n = 119 cells; NEMOer-m, n = 117 cells; NEMOer-s, n = 106 cells NEMOer-f, n = 143 cells; NEMOer- c, n = 134 cells.). Typical cells showing ER Ca^2+^ oscillations recorded in bath solution containing 2 mM Ca^2+^ (C). n = 3 independent biological replicates, with at least 17 cells per repeat. Data shown as mean ± s.e.m. (D) Simultaneous monitoring of 10 μM CCh-induced Ca^2+^ oscillations within the ER and cytosol using NEMOer-f and a cytosolic red indicator, R-GECO1.2 in HEK293 cells. Both GECIs are linked together by ER-membrane-spanning domain of STIM1 and expressed at a 1:1 ratio, with NEMOer-f and R-GECO1.2 facing ER lumen and cytosol, respectively. Left, representative traces; right, enlargement of one typical oscillating pulse (3 independent biological replicates). (E-G) Ratiometric signals of m<u>Tu</u>rquoise2- <u>N</u>EMO<u>er</u>-m (TuNer-m) transiently expressed in COS-7 cells with multimodality structured illumination microscopy (Multi-SIM). The design of TuNer is shown in (**Extended Data Fig. 3A**). The ratio (R) of TuNer indicators is defined as R = F_NEMOer_/F_sfTq2ox_. (E) Typical basal fluorescence or ratiometric images (scale bar, 10 μm), with an enlarged portion of the cell shown in the bottom row (scale bar, 5 μm). Regions indicated by purple arrows show higher fluorescence signal but lower TuNer-m ratio, as compared with their corresponding adjacent areas. Those marked by yellow arrows show higher TuNer-m ratio but lower NEMOer-m fluorescence as compared to their adjacent areas. (F-G) Comparison of TuNer-m ratios within ER sheets and ER tubules. (F) Diagram showing the selection of Region of Interest (ROI) in ratio image: ER sheets (white circles), edge of ER sheets (yellow circles) or ER tubules (green circles) (G) Statistics of TuNer-m ratio within areas illustrated in (F) (**** *P*<0.0001, * *P*=0.0267, one-way ANOVA, n=76 ROIs from at least 5 cells).

To enable monitoring of ER Ca^2+^ homeostasis, we generated ratiometric m<u>Tu</u>rquoise2-<u>N</u>EMO<u>er</u> (TuNer) indicators by using a mTurquoise2 (mTq2) variant as both an indicator for the sensor expression level and a Föster energy resonance transfer (FRET) donor for NEMOer sensors (**Extended Data Fig. 3A**). We reasoned that the FRET efficacy between mTq2 and NEMOer indicators may be proportional to NEMOer fluorescence. This would ensure reciprocal change in NEMOer and mTq2 fluorescence, resulting in amplified NEMOer responses (**Extended Data Fig. 3B**). Since ER Ca^2+^ level is less conveniently manipulatable as compared with cytosolic Ca^2+^, we used cytosolic mNeonGreen based indicator NCaMP7 ^26^ to optimize the linkers and interfaces between the FRET pair (**Supplementary Table 4**), and then applied the optimized strategy for NEMO and NEMOer sensors to obtain ratiometric m<u>Tu</u>rquoise2–<u>N</u>EMO (TurN, cytosolic version) or TuNer sensors (**Supplementary Table 5&6**). The DRs of all these TurN indicators were 1.4∼2 fold higher than the corresponding mono-fluorescent templates (**Supplementary Table 4∼6**). When stimulated with 10 μM CCh, decreases in ratios of TuNer indicators were more than 1.5 fold of miGer, a ratiometric ER Ca^2+^ indicator based on G-CEPIA1er (**Extended Data Fig. 3C**). When measured with TuNer sensors, the basal ER Ca^2+^ levels within HEK293 and HeLa cells fell in the range of 0.74-1.55 mM (**Extended Data Fig. 3D**).

We further examined TuNer-m responses with a commercial confocal imaging system. Tubular ER structure within COS-7 cells transiently expressing TuNer-m was clearly visible with good signal-to-noise ratio (**Extended Data Fig. 4A&4B**). Nuclear Ca^2+^ signaling is crucial for proper cell function^27^, yet nuclear ER Ca^2+^ content were less studied. Here we observed that basal nuclear ER Ca^2+^ levels are slightly, but significantly higher than the rest portion of ER (indicated by white and yellow arrows, respectively, **Extended Data Fig. 4A&4C**). Further analysis showed that the TuNer-m ratio also differed significantly among different subcellular areas at rest, demonstrating uneven distribution of resting ER Ca^2+^ levels (**Extended Data Fig. 4D**). When examined with structured illumination microscopy (SIM) super-resolution imaging, the TuNer-m ratios within ER tubules and on the edge of ER sheets are higher than the majority of ER sheets in COS-7 cells (**Fig. 2E-2G**), suggesting that Ca^2+^ levels within ER sheets are significantly lower than those in their corresponding neighboring ER tubules.

Of note, the basal subcellular NEMOer-m fluorescence does not correlate with that of TuNer-m ratio (microdomains indicated by arrows, **Fig. 2E** and **Extended Data Fig. 4B**), clearly demonstrating an uneven distribution of GECIs within ER. Upon stimulation with 5 μM ATP, different subcellular regions of COS-7 cells showed repetitive fluctuations of TuNer-m ratios with varying amplitudes, illustrating slightly unsynchronized Ca^2+^ oscillations (**Extended Data Fig. 4E, Supplementary Video 1**). The discrepancies between NEMOer-m signals and TuNer-m ratios within microdomains are more pronounced in enlarged pictures or zoom-in videos (Dots indicated by white arrows, **Extended Data Fig. 4B and Supplementary Video 2**). These results thus underscore the need for caution when interpreting results obtained with intensiometric GECIs.

During ER Ca^2+^ oscillations induced by 5 μM ATP (**Extended Data Fig. 4E**) as well as dynamic re-arrangements of ER tubules, we could clearly see transient punctate microdomains with higher TuNer-m ratios (Dots indicated by white arrows, **Supplementary Video 3**), highlighting the highly discrete nature of Ca^2+^ refilling within ER tubules. Transient “Moving dark dots” with decreased TuNer-m ratios were also observed (pointed by yellow arrows, **Supplementary Video 3**), likely representing the brief reductions of ER Ca^2+^ levels within microdomains of ER tubules that are responsible for propagating cytosolic Ca^2+^ waves.

Collectively, our data show that the bright and sensitive TuNer sensors are well suited for super-resolution imaging of dynamic ER Ca^2+^ signaling in non-excitable cells.

## Performance of NEMOer sensors in cultured neurons

We proceeded to investigate NEMOer responses in cultured primary hippocampal neurons under electrical field stimulation. Since it is established that excited neurons often show ER Ca^2+^ overload ^14,24,28^, we selected a NEMOer variant with a much lower affinity (K_d_=19.2±4.0 mM), designated it as NEMOer for neurons (NEMOer-n), and examined its ability to report ER Ca^2+^ signals induced by electrical field stimulation. In transiently transfected neurons, NEMOer-n fluorescence displayed an ER-like reticular network resembling the endoplasmic reticulum in both dendrites and soma. With a train of 20 action potentials (APs) at 10 Hz stimulation, we observed a robust increase in NEMOer-n fluorescence, with dendrites showing a more pronounced response compared to the soma (**Fig. 3A-B**), indicating more ER Ca^2+^ overload in dendrites. Subsequently, we detected Ca^2+^ signals elicited by a single AP in secondary dendrites of hippocampal neurons. Compared to signals in G-CEPIA1er-expressing neurons, the single-AP-induced transient Ca^2+^ response in NEMOer-n-expressing neurons was significantly enhanced (**Fig. 3C**), while the other NEMOer variants showed a similar response.

**Figure 3.**
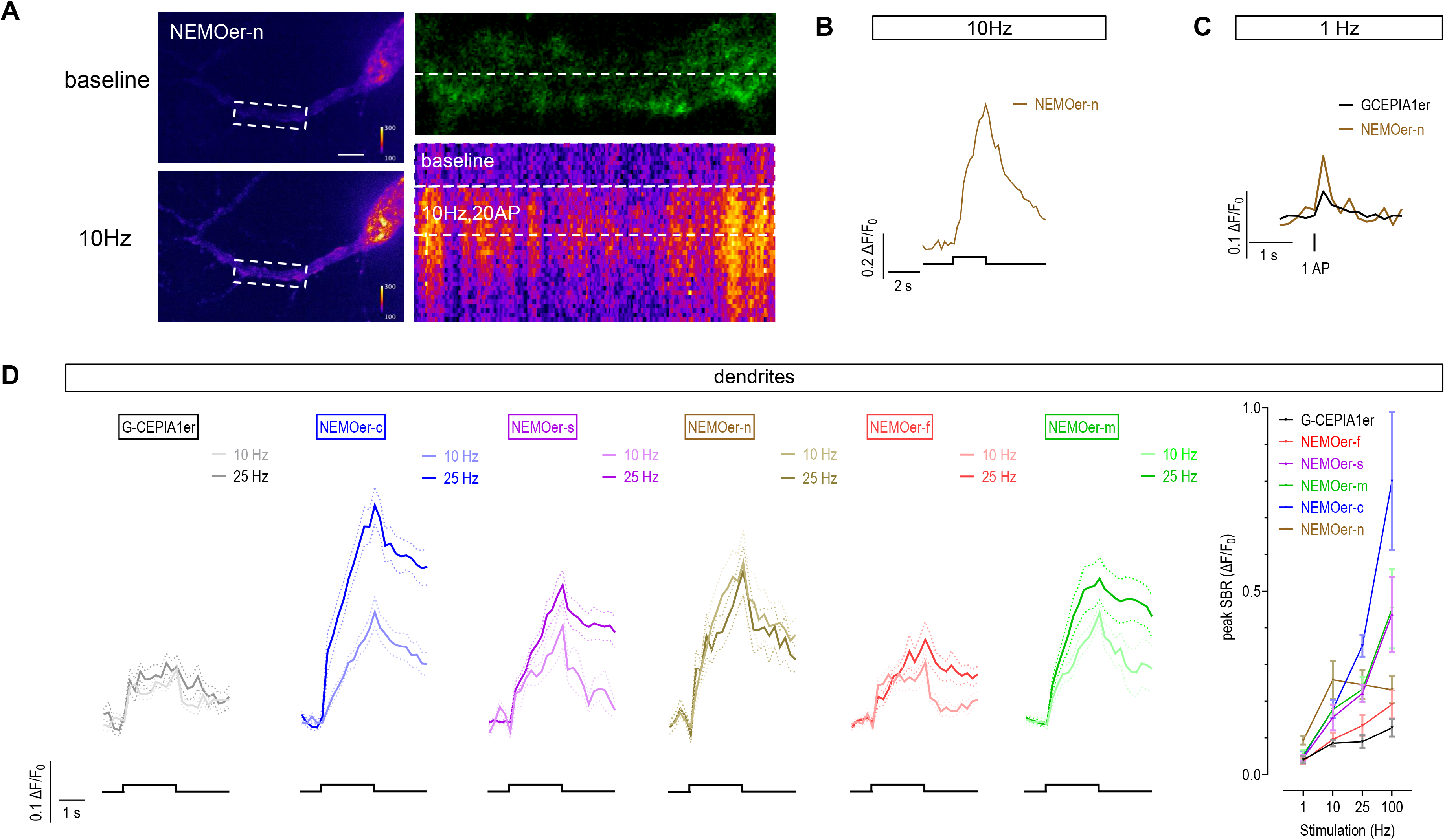
Responses of NEMOer variants or G-CEPIA1er to electric field stimulation in rat hippocampal neurons. (A) Typical images showing the fluorescence distribution and responses of NEMOer-n to 10 Hz field stimulation. Pseudocolor cellular images of NEMOer-n before (top left) and after 20 AP (10 Hz) stimulation (bottom left) (scale bar, 10 μm). Enlarged view of the dendrite region (top right), with a white dashed line indicating the position of line scan. Kymograph showing NEMOer-n responses to 10 Hz stimulation within a dendritic region (bottom right) along the indicated line. Scale bar, 10 μm. (B) Representative tracing of NEMOer-n responses to 10 Hz stimulation. (C) Typical response of NEMOer-n or G-CEPIA1er to 1 Hz stimulation. (D) Response of NEMOer variants or G-CEPIA1er within dendrites induced by electric stimulation with varies frequencies, with mean response curves shown on the left and statistics shown on the rightmost. (For stimulation at varied frequencies, G-CEPIA1er, n = 12,11,12,14 cells; NEMOer-f, n = 15, 14, 12, 13 cells; NEMOer-s, n = 11, 13, 12, 13 cells; NEMOer-m, n = 14, 17, 16, 13 cells; NEMOer-c, n = 13, 14, 12, 12 cells; NEMOer-n, n = 12, 15, 14, 13 cells.).

Next, we examined the responses of other NEMOer variants following stimulations of 10 Hz, 25 Hz or higher frequency, and found that NEMOer sensors had a dramatic increase in the peak signal-to-background ratio (SBR) compared to the G-CEPIA1er sensor. Notably, the NEMOer-c sensors had the highest response amplitude, while NEMOer-n and NEMOer-f was fast enough to pick up individual responses to 10 Hz stimulation (**Fig. 3C-D**). To further verify if the responses reported by NEMOer sensors represent Ca^2+^ increase in ER, we examined the effects of SERCA inhibition by incubating cells with cyclopiazonic acid (CPA) for one hour and then stimulating them at a frequency of 100 Hz for 2 seconds. As expected, the NEMOer-c and NEMOer-s signals were significantly reduced in response to the stimulation in the presence of CPA compared to DMSO (0.33 vs 0.81 (peak response in CPA vs DMSO), *P*=0.02; 0.12 vs 0.46 (peak response in CPA vs DMSO), *P*=0.004, respectively). These results suggest that NEMOer sensors have a significantly superior ER Ca^2+^ responses to physiological frequency stimulations in primary neurons compared to G-CEPIA1er sensors.

## Performance of NEMOer sensors in cultured cardiac muscle cells

We moved on to further compare the responses NEMOer sensors with G-CEPIA1er in neonatal cardiomyocytes challenged with 30 mM caffeine, an agonist of ryanodine receptors (RyR), the primary Ca^2+^ releasing channels on SR. Decreases in NEMOer signals were more than 80%, significantly larger than those of G-CEPIA1er or ER-GCaMP6-150 (∼66%). Specifically, NEMOer-f demonstrated the fastest kinetics among NEMOer variants (**Fig. 4A**), significantly faster than ER-GCaMP6-150 (decay rate constant: 2.47±0.06 sec^-1^ vs 0.81±0.02 sec^-1^, *P* < 0.0001), indicating its capability to detect rapid ER Ca^2+^ dynamics in cardiac muscle cells.

**Figure 4.**
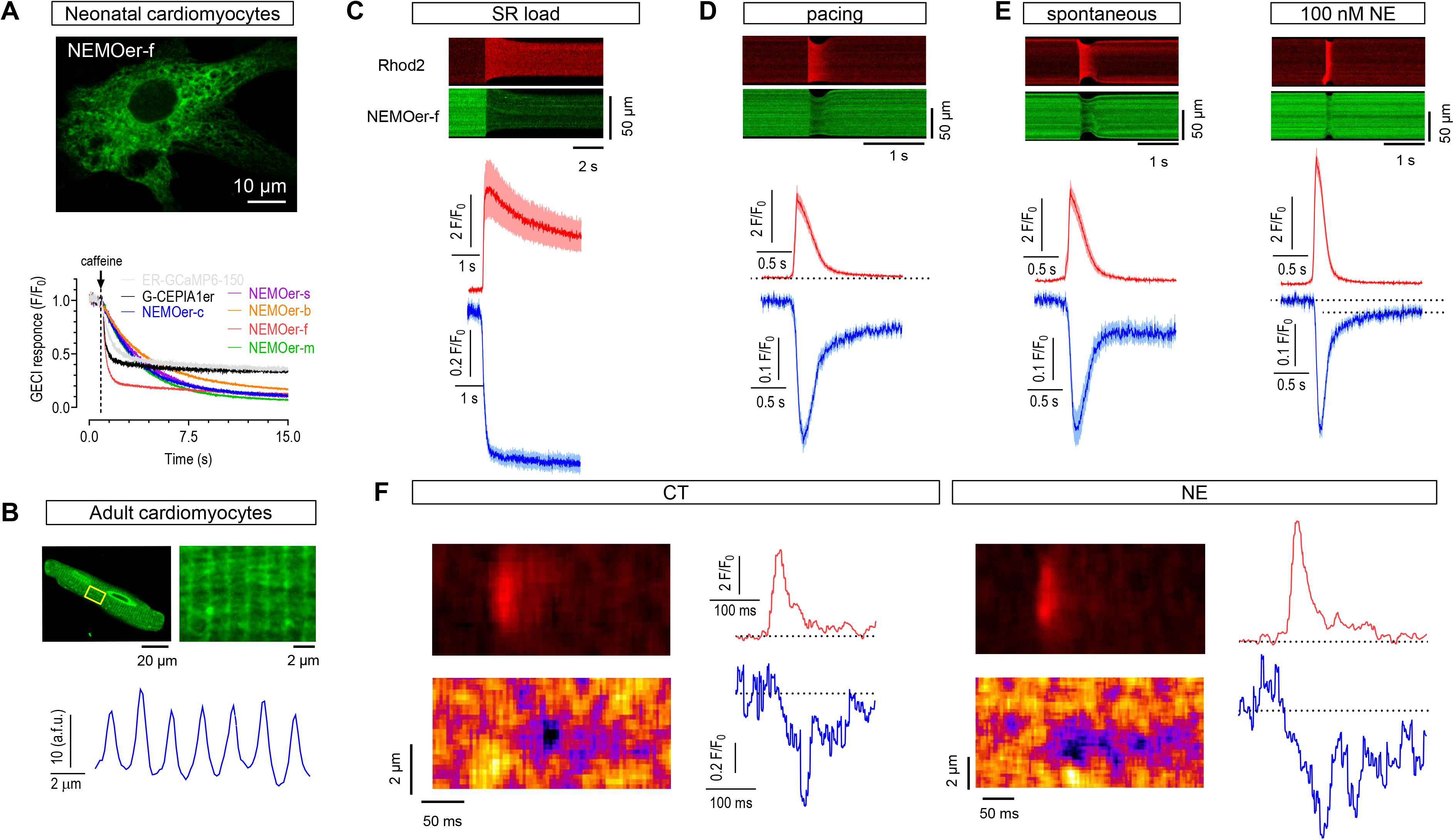
Performances of NEMOer sensors in monitoring SR Ca^2+^ dynamics in cardiomyocytes. (A) SR load measurement by 30 mM caffeine perfusion in neonatal rat cardiomyocytes. Top, pseudocolor image showing a typical cell expressing NEMOer-f; bottom, averaged time curves showing caffeine-induced responses of transiently expressed indicators. Arrow indicated the time caffeine stimulation. Scale bar, 10 μm. (B) Subcellular distribution of NEMOer-f fluorescence in adult rat cardiomyocyte revealed with confocal imaging. Top left, typical confocal images (scale bar, 20 μm); top right, enlarged view of boxed area shown in top left image; bottom trace, spatial profile for the boxed region. (C-E) Local and global cytosolic and SR Ca^2+^ dynamics in adult rat cardiac myocytes revealed by dual-color imaging using NEMOer-f and a cytosolic Ca^2+^ indicator, Rhod-2. Top, typical line-scan images; bottom, corresponding mean time course plots. Ca^2+^ responses induced by caffeine (30 mM) (C). Local SR Ca^2+^ scraps and accompanying cytosolic Ca^2+^ transient trigged by electric field stimulation (D) or during a spontaneous release (E), either under control (CT) (left) or when treated with 100 nM NE (right) (For each trace, n = 6-18 cells from 3 adult rats). (F) Spontaneous elementary cytosolic Ca^2+^ spark and SR Ca^2+^ blink events under CT (left) or NE-treated (right) conditions revealed by Rhod-2 and NEMOer-f, respectively. Typical line-scan images are shown on the left; corresponding time-course curves are shown on the right.

We then infected adult rat cardiomyocytes with adenovirus carrying NEMOer-f and imaged them with confocal microscopy 36 ∼ 48 hours later. We found that NEMOer-f fluorescence exhibited a striated sarcomeric pattern, indicating its correct localization in the SR (**Fig. 4B**). To better assess NEMOer-f performance, we co-loaded cardiomyocytes with a cytosolic chemical Ca^2+^ indicator Rhod-2, and simultaneously monitored the responses of both indicators. In response to 30 mM caffeine, Rhod-2 signal increased approximately four-fold, indicating a substantial drop in SR Ca^2+^ levels. Concurrently, NEMOer-f fluorescence rapidly decreased to about 75% of baseline, significantly surpassing values reported with a chemical indicator Fluo-5N (66.7%) ^29^. This result highlights the high sensitivity of NEMOer-f for directly monitoring SR Ca^2+^ decreases in intact cardiomyocytes (**Fig. 4C**).

Next, we examined Ca^2+^ dynamics during excitation-contraction (E-C) coupling with NEMOer-f and Rhod-2. In response to field electrical stimulation (15 V, 1 Hz), cytosolic Ca^2+^ pulse indicated by Rhod-2 signal (ΔF/F_0_ = 3.1 ± 0.3) was accompanied by a large transient decrease in NEMOer-f fluorescence (ΔF/F_0_ = 0.35 ± 0.02) (**Fig. 4D, Extended Data Fig. 5A**), demonstrating simultaneous lowering of local SR Ca^2+^ levels, or Ca^2+^ scraps. Compared with those reported by a commercial Ca^2+^ dye (Fluo5N, ΔF/F_0_ = 0.1 ∼ 0.2) or R-CEPIA1er ^30,31^, Ca^2+^ scarps indicated by NEMOer-f demonstrated a similar time-to-nadir (126.8 ± 11.4 ms) and recovery time (414.8 ± 26.9 ms). Nevertheless, peak NEMOer-f responses was larger than R-CEPIA1er or Fluo-5N, showcasing its increased sensitivity over these two sensors.

Using Rhod-2 and NEMOer-f, we subsequently examined the corresponding spontaneous cytosolic and SR Ca^2+^ activities under resting conditions or upon adrenergic receptor activation (100 nM norepinephrine, NE). We found that both Rhod-2 and NEMOer-f responses were similar to those triggered by field stimulation (**Fig. 4E, left**). In line with previous reports^31^, cardiomyocytes under adrenergic activation exhibited significantly larger spontaneous Rhod-2 transients, concomitant with faster responses from NEMOer-f (**Fig. 4E, right; Extended Data Fig. 5B&C**).

We further assessed the ability of NEMOer-f to detect Ca^2+^ blinks, the elementary decreases in SR Ca^2+^ level leading to Ca^2+^ sparks, the fundamental units of excitation-contraction coupling in cardiomyocytes. We successfully observed the Ca^2+^ spark-blink pairs by Rhod-2 and NEMOer-f, respectively (**Fig. 4F)**. The peak amplitude of Ca^2+^ blink indicated by NEMOer-f (ΔF/F_0_ = 0.79 ± 0.02), around three folds of the Fluo-5N signal documented in rabbit muscle cells ^22,32^. The duration of blink response indicated by NEMOer-f was slower than those reported by Fluo-5N. Further researches are needed to determine if the observed effect is due to slower speed of NEMOer-f or facilitated Ca^2+^ diffusion by Fluo-5N, a smaller Ca^2+^ buffer molecule within SR. Nevertheless, to the best of our knowledge, this marks the first measurement of Ca^2+^ blinks with genetically encoded Ca^2+^ sensors, underscoring NEMOer-f’s exceptional sensitivity.

## *In vivo* performance of NEMOer indicator in zebrafish

To test NEMOer-f *in vivo*, we took the advantage of larval zebrafish that is optically transparent for live imaging. We constructed Tol2 transposase-based transgenic vectors that express codon-optimized NEMOer-f specifically in the SR of muscle cells using the *β*-actin promoter ^33^. By injecting the fertilized eggs at their 1-cell stages, we obtained larvae with transient expression of NEMOer-f and imaged sparsely labeled muscle cells on day 5 post fertilization (dpf) (**Fig. 5A**). As expected, the baseline fluorescence of NEMOer-f indicated the SR localization of the sensor in muscle cells with a typical striated morphology (**Fig. 5B**). When muscle contracted, NEMOer-f fluorescence exhibited robust signals with a sharp decrease followed by a slow rising phase (**Fig. 5C-D**, **Supplementary Video 4**). The median responses of NEMOer-f signals are around 30% (**Fig. 5E**). These data demonstrate that NEMOer-f functions in intact animals to detect spontaneous Ca^2+^ releases in SR.

**Figure 5.**
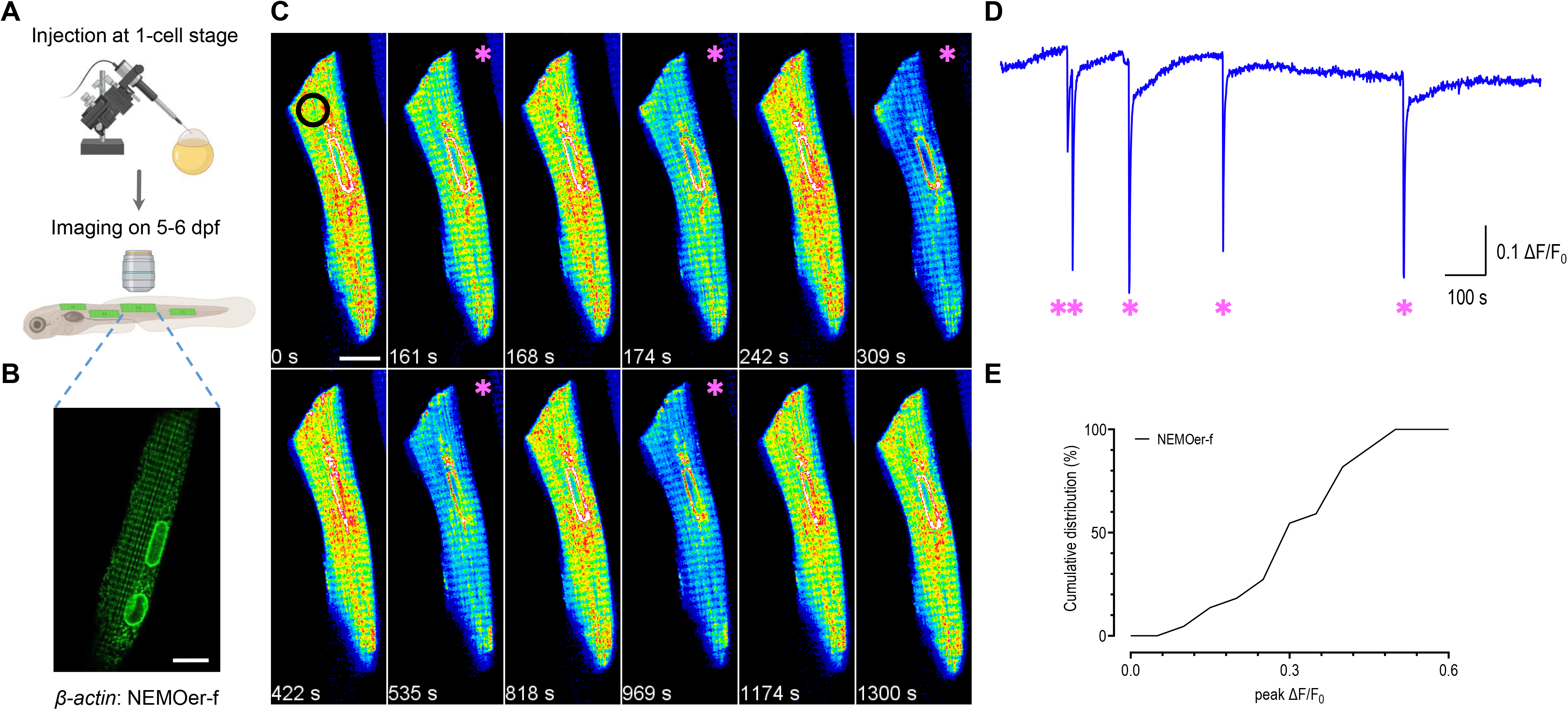
*In vivo* NEMOer-f responses to spontaneous SR Ca^2+^ releasing events in skeletal muscles of zebrafish. (A) A schematic depiction of the NEMOer-f *in vivo* imaging workflow in zebrafish. (B) A typical sparsely labeled skeletal muscle cell, expressing NEMOer-f. Scale bar, 10 μm. (C) Image montage of time-lapse imaging at 1 Hz. Black circle indicates the ROI for extracting time series in (D). Pink asterisks indicate single Ca^2+^ events. Scale bar, 10 μm. (D) Representative time series showing spontaneous SR Ca^2+^ release indicated by NEMOer-f signal. Pink asterisks indicate single Ca^2+^ events. (E) Cumulative distribution of amplitude NEMOer-f responses (n = 22 events from 4 zebrafish larvae).

## Conclusions

Here we reported NEMOer, a highly sensitive green GECIs for ER/SR with large dynamics. In comparison to commonly used GECIs derived from GCaMP-like series, such as G-CEPIA1er, NEMOer indicators show 10-fold higher brightness with the dynamic ranges 50 times larger, enabling the detection of physiological ER/SR Ca^2+^ dynamics with significantly larger signals. NEMOer sensors also exhibit significantly enhanced photochemical properties, showing minimal sensitivity to physiological pH variations and enduring 50 times more illumination without apparent bleaching. These ideal properties make NEMOer compatible with super-resolution recording of SR/ER Ca^2+^ homeostasis and dynamics. Remarkably, the fast NEMOer variant, NEMOer-f, stands out as the inaugural GECI enables the detection of Ca^2+^ blink, the elementary Ca^2+^ releasing signal from SR of cardiomyocytes, as well as *in vivo* detection of spontaneous SR Ca^2+^ releasing event in a model organism zebrafish. Collectively, the exceptionally dynamic NEMOer sensors emerge as the premier choice for monitoring Ca^2+^ dynamics and homeostasis in mammalian cells, particularly within muscle cells.

## ACKNOWLEDGMENTS

This work was supported by the National Natural Science Foundation of China (92254301 and 91954205 to Y. W., 32171026 to Y. M., 32250003 and 32230048 to S.-Q.W.), STI2030-Major Projects (2021ZD0202503 to A.-H. T., 2021ZD0204500 and 2021ZD0203704 to Y. M.), the National Key Research and Development Program of China (2020YFA0112200 to A.-H. T.), China Postdoctoral Science Foundation General Program (2022M723246 to Z. W.), and Scientific Instrument Developing Project of the Chinese Academy of Sciences (YJKYYQ20210029 to Y. M.) and the Ministry of Science and Technology of China (2019YFA0802104 to Y. W.).

## Author contributions

Y. W., A-H. T., S-Q W, and Y. M. supervised and coordinated the study. W. G. and J. L. designed and generated all the plasmid constructs, with help from Z.Z.. J. L., W. G. and H. Z. performed the *in vitro* assays. W. G. performed all fluorescence imaging in non-excitable cells, with help from J. L.. J. C. performed confocal imaging of dissociated neurons. Y. Z. performed confocal imaging of dissociated cardiac cells. W. G., A. J., and H. Z. performed Multi-SIM imaging on COS-7 cells under the guidance of Y. W. and D. L.. Z.W. performed *in vivo* Ca^2+^ imaging of muscle cells zebrafish with help from S. W.. C. X. provided tech support for live cell imaging. W. G., J. C., Y. Z., Z. W., J. L. and H. Z. analyzed data, with input from the other authors. D. L. G., S. W., Y. M., T. H., and J. L. provided intellectual input to the manuscript. Y. M., A-H. T. and Y. W. wrote the manuscript with inputs from all the other authors.

## Acknowledgments

We would like to thank the Experimental Technology Center for Lifesciences, Beijing Normal University.

---

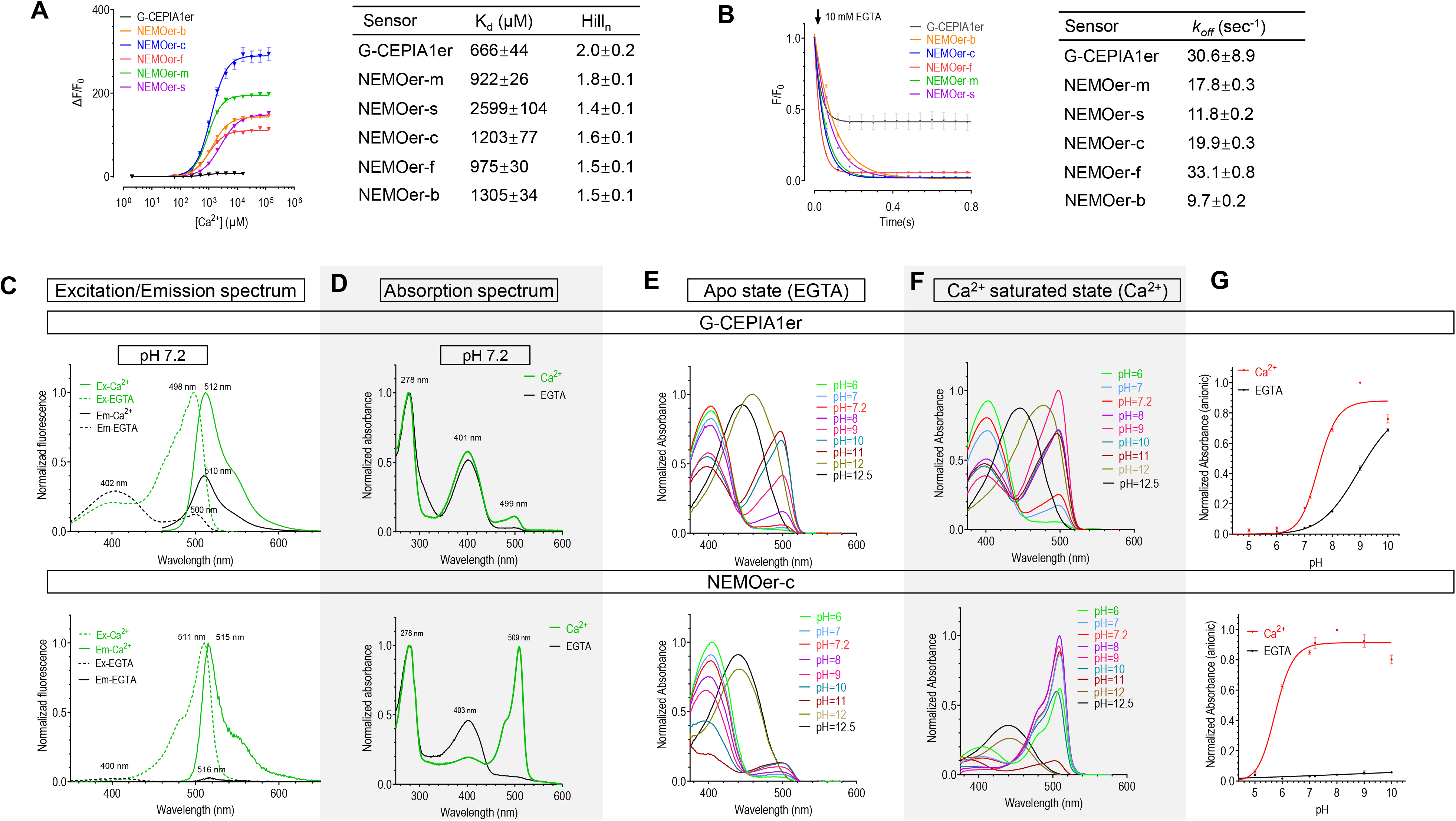

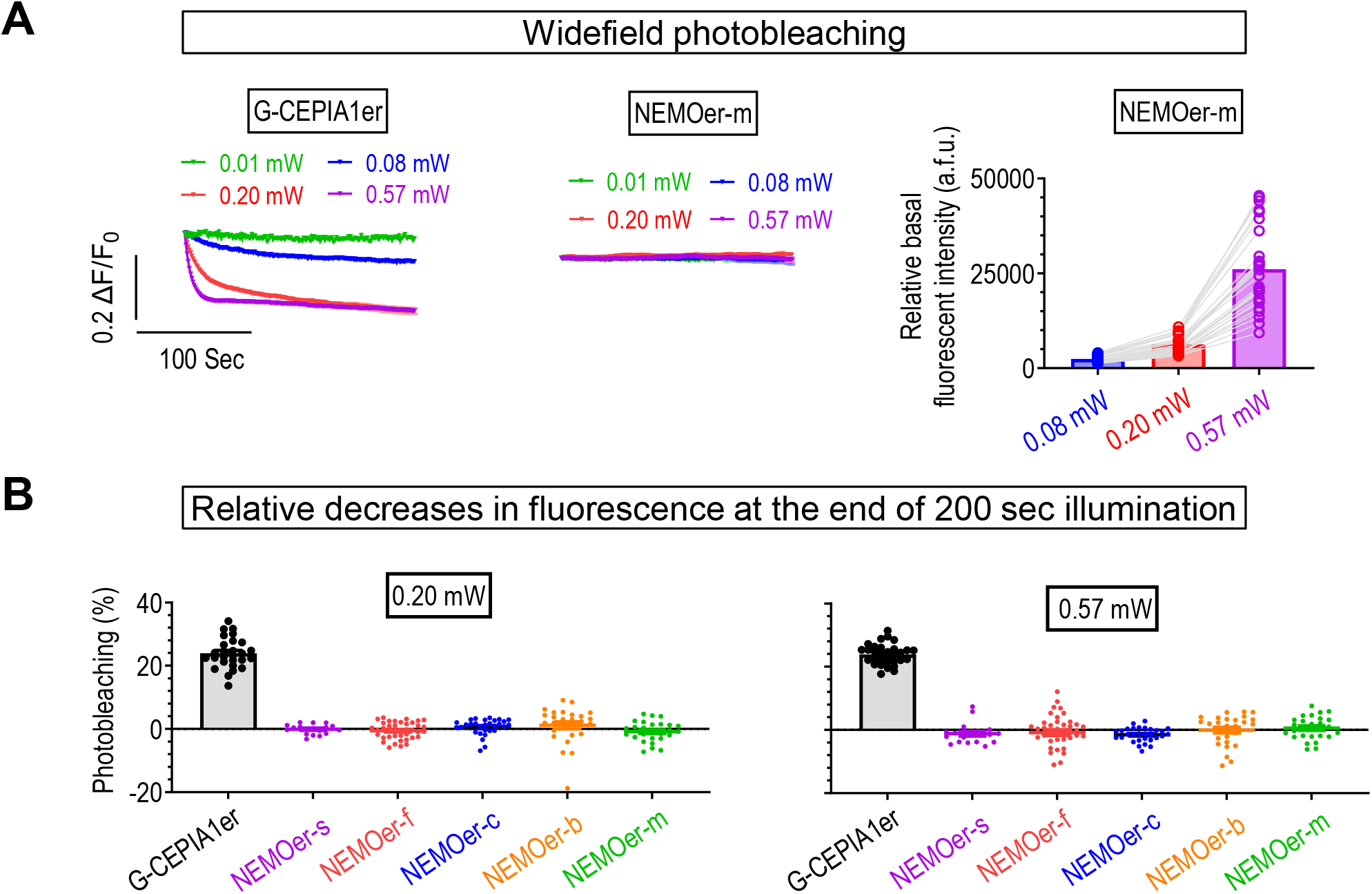

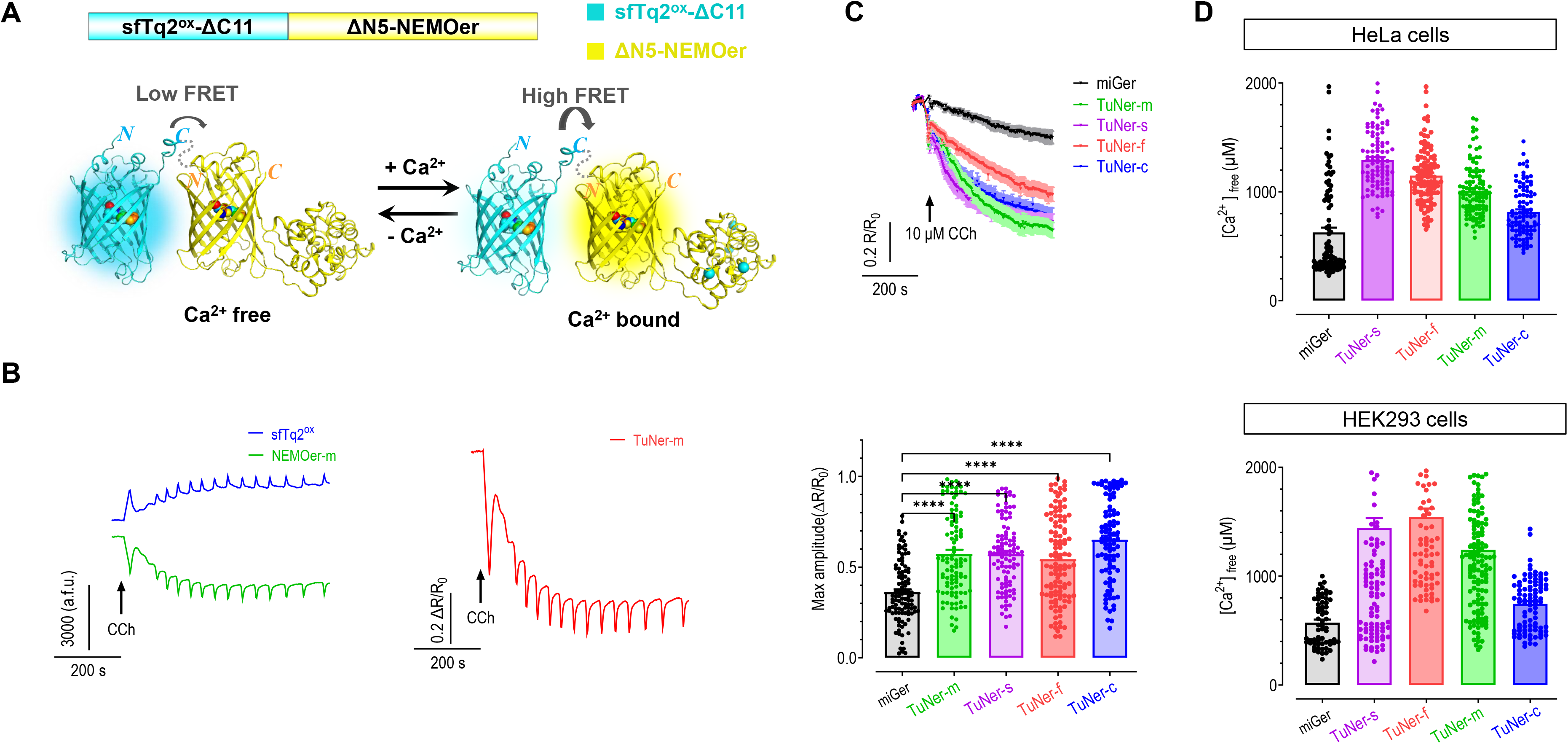

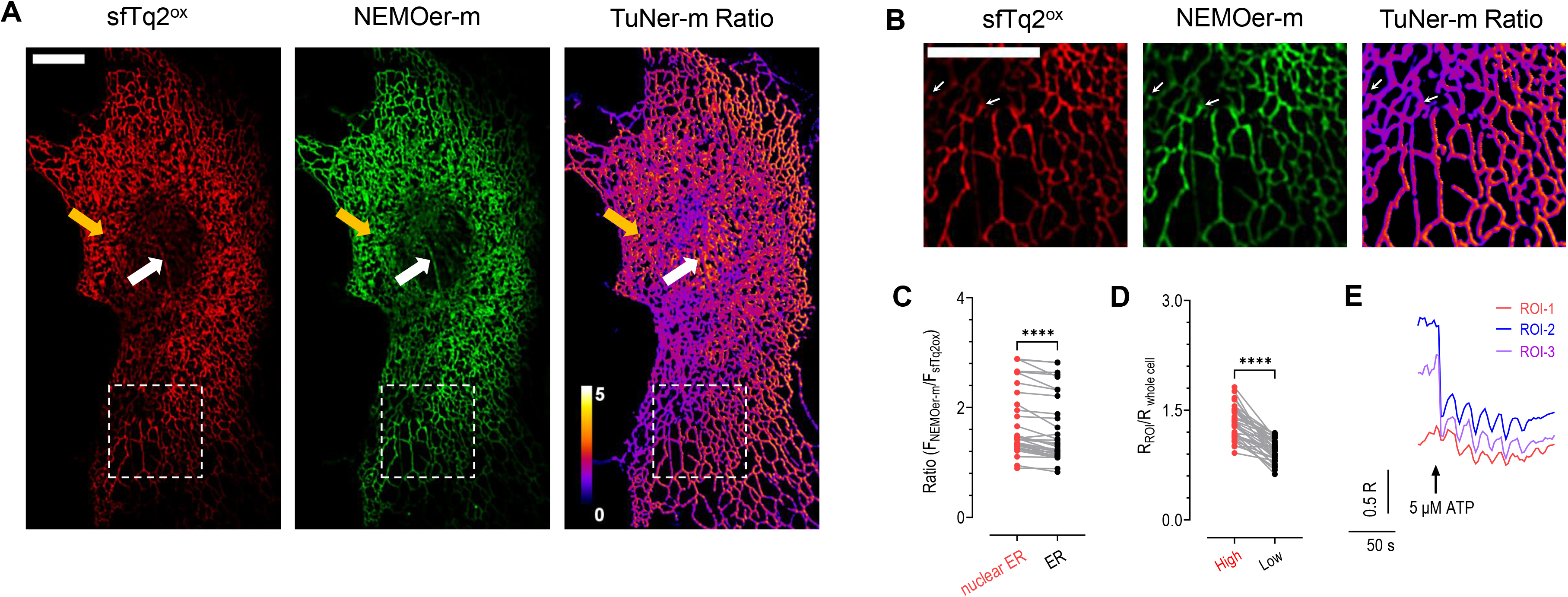

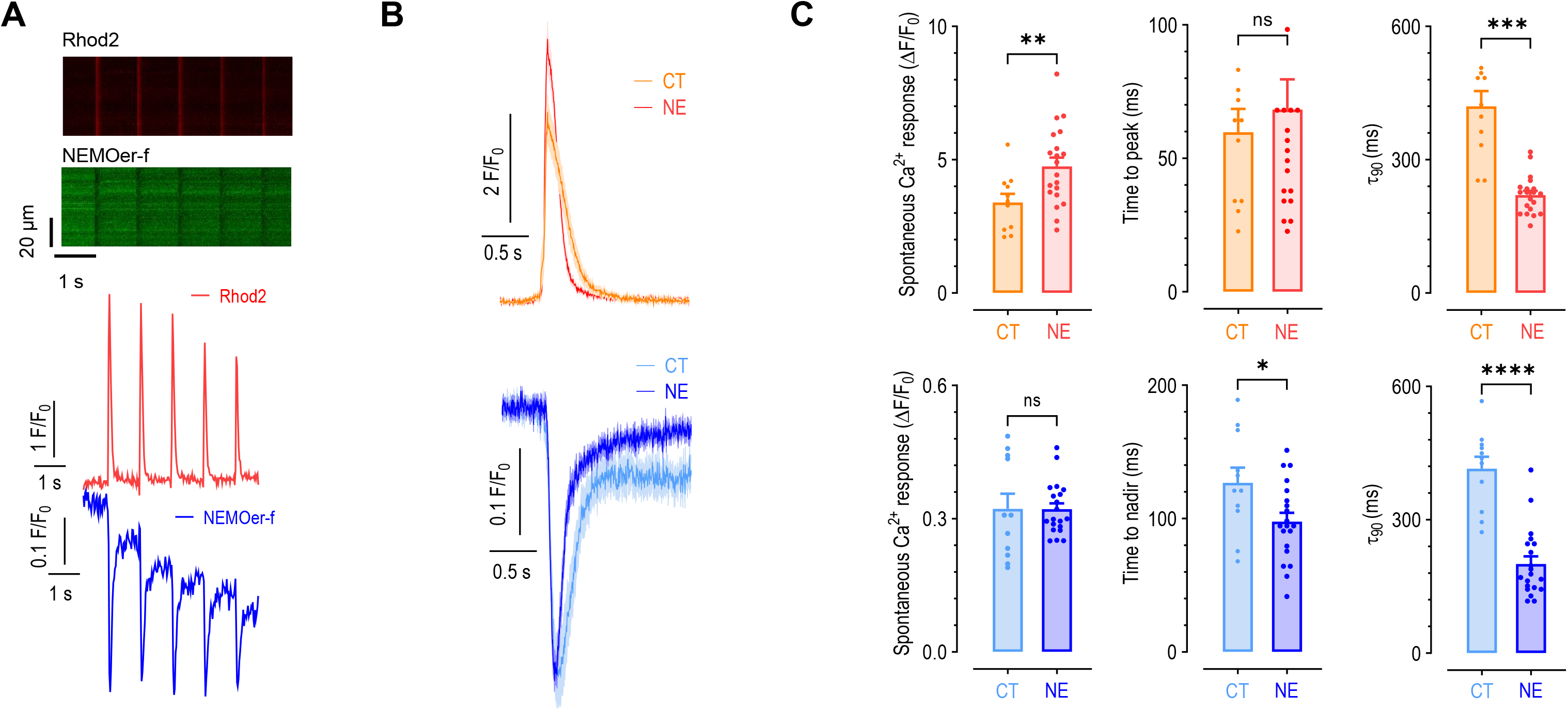

