## Supplementary Table 1-7 for "Highly Dynamic and Sensitive NEMOer Calcium Indicators for Imaging ER Calcium Signals in Excitable Cells"

**Supporting Information (SI)**

**Methods**

***Plasmids construction and Recombinant Adenovirus Production***

ER-GCaMP6-150 and superfolder turquoise2^ox^ (sfTq2^ox^)^1^ was synthesized and incorporated into a pCDNA3.1(+) vector by BGI Geneland Scientific Co., Ltd, Jiangsu, China. Zebrafish codon-optimized NEMOer-f was synthesized by the entrusted company, Beijing Tsingke Biotech Co., Ltd., Beijing, China. The construction of other plasmids followed previously described methods. Briefly, target fragments were PCR amplified from corresponding templates, and the vector backbone was linearized with appropriate restriction enzymes. Corresponding mutations or substitutions were all included in primers. Fragments were then reassembled and inserted into the linearized vector utilizing a Ready-to-Use Seamless Cloning Kit (B632219, Sangon Biotech, Shanghai, China). Plasmids were all confirmed by sequencing. The vector backbones were linearized using the following restriction enzymes: BamHI and EcoRI for pCDNA3.1, EcoRI and XhoI for pTol2-β-actin, and NcoI and XhoI for pET28a.

To construct variant library of NEMOer candidates, we performed PCR amplification of NEMOs^2^, incorporating known Ca^2+^-affinity-reducing mutations^3^ via primers. Subsequently, we added the endoplasmic reticulum localization signal sequence from GCEPIA1er^3^ and the coding sequence for ER-resident sequence KDEL to the N and C-termini of the constructs, respectively. Then, we reassembled and inserted them into a pCDNA3.1 vector backbone. To construct the TurN (mTurquoise2–NEMO)/TuNer (mTurquoise2–NEMOer) plasmids, we amplified the coding sequence of sfTq2^ox^-ΔC11 from pCDNA3.1(+)-sfTq2^ox^ and the coding sequences of NEMO/NEMOer-ΔN5 from NEMO/NEMOer, respectively. Subsequently, we reassembled and inserted them into a custom-made pCDNA3.1(+) vector with puromycin resistance. To generate the mKate-P2A-GECI plasmids, we assembled the coding sequence of mKate-P2A, which was amplified from the mKate-P2A-NEMO plasmid^2^, along with the coding sequences of NEMOer/GCEPIA1er, into the custom-made pCDNA3.1 vector. To create ER-membrane anchoring GECI with NEMOer-f and RGECO1.2 facing ER lumen and cytosol, respectively, we utilized LIMETER(Addgene #113933)^4^ as the backbone, PCR amplified the coding sequence of STIM1 signal peptide, NEMOer-f, ER-membrane-spanning domain of STIM1 (STIM1_195-240_), and R-GECO1.2 (Addgene #45494)^5^. These sequences were recombined into the custom-made pCDNA3.1 vector using G, GGRSGG and (GA)_6_ linkers, respectively. For expression in *Danio rerio*, codon-optimized NEMOer-f was subcloned into the pTol2-β-actin vector, resulting in the generation of pTol2-β-actin: NEMOer-f.

The NEMOer-f gene sequences was amplified and inserted into pENTR2B vector (Invitrogen). The adenovirus was produced using the adenoviral expression system (Invitrogen) in HEK293A cells.

***Bacterial expression and protein purification***

Transetta (DE3) bacteria (TransGen Biotech, Beijing, China) were transformed with pET28a plasmids carrying GECI coding sequences and cultured in LB medium until reaching an optical density (OD) of 0.5. Protein expression was then induced overnight at 20 °C by the adding 300 mM isopropyl-β-d-thiogalactoside (IPTG). The recombination proteins were purified following a previously described protocol^6^. Briefly, cells were harvested by centrifugation and re-suspended in 20 mL buffer 1 (mM, 20 Tris, 300 NaCl and 1 imidazole, pH 7.2). Cells were initially lysed and subsequently subjected to ultrasonic disruption, followed by incubation with 1 mL Ni Sepharose (17-5318-01, GE Healthcare, Piscataway, NJ, USA) to enrich recombinant GECI proteins with histidine-tags. The columns were sequentially washed with 20 mL buffer 1 and 10 mL buffer 2 (mM, 20 Tris, 500 NaCl and 10 imidazole, pH 7.2). Finally, the GECI proteins were eluted using 5 mL buffer 3 (mM, 20 Tris, 100 NaCl and 300 imidazole, pH 7.2).

***In vitro Ca^2+^ titrations and kinetics measurements***

The fluorescence intensity from buffers solutions containing 50 μg/mL GECI proteins, used to calculate Ca^2+^ dissociation constant (K_d_), were recorded by a multi-mode microplate reader (Flexstation 3, Molecular Devices, USA). The buffer solution contained 130 mM KCl, 50 mM MOPs ^7^ and CaCl_2_ with varying concentrations ranging from 0 mM to 128 mM (pH 7.2). The excitation/emission wavelengths of GCEPIA1er and NEMOer were 485 /510 nm and 490 / 520 nm with 10 nm bandpass, respectively. Ca^2+^ titration curves of Ca^2+^ concentration versus relative fluorescence intensity were then plotted and fit with specific binding with hill slope function^2^ in Prism 9.5 software.

The fluorescence traces used to calculate disassociation kinetics (*k_off_*) were recorded by a fluorescence polarization microplate reader (POLARstar Omega, BMG LABTECH, Germany). Fluorescence baselines from buffers containing 50 μg/mL GECI proteins, 5 mM Ca^2+^, 50 mM MOPS, 100 mM KCl, pH 7.2 ^8^ were first recorded. Subsequently, changes in fluorescence were then monitored in response to rapidly mixing with a buffer containing 10 mM EGTA, 50 mM MOPS, 100 mM KCl, pH 7.2. The excitation/emission wavelength of GCEPIA1er and NEMOer was 485/520 nm. The *k_off_* value was determined by fitting the fluorescence decay curve with a single exponential function using Prism 9.5 software.

***Spectra recording and measurement of quantum yields of GECIs***

Purified proteins were diluted to a final concentration of 50~100 μg/mL in a buffer containing 130 mM KCl, 50 mM MOPs, supplement with or without 128 mM CaCl_2_, pH 7.2. Fluorescence spectra were measured with a FS5 spectrophotometer (Edinburgh Instrument, Scotland) controlled by Fluoracle software. Absorption spectra were measured with an ultraviolet and visible spectrophotometer (UV2600, Shimadzu, Japan) controlled by UVprobe software.

The emission spectra and optical density(OD) used to calculate Quantum yields (Φ) of GECIs were determined with a FS5 spectrophotometer and a an ultraviolet and visible spectrophotometer ^2^. We used the obtained the emission spectra to calculate the total integrated fluorescence intensity with different concentrations of GECI protein. Linear regression of integrated fluorescence intensity and absorbance values to get the slopes (S). Φ was determined by the equation: Φ_protein_ = Φ_standard_ × (S_protein_/S_standard_)^9^. The reference standards are fluorescein (φ=0.925) and TOLLES (φ=0.79) in 0.1 M NaOH ^10^, excited at 470 nm and 405nm, respectively^11^.

***Chromophore extinction coefficients***

The absorption spectra of buffer solutions containing 200 μg/ml GECI proteins, used to calculate extinction coefficients (ε), were recorded by Flexstation 3. Buffer solutions containing 30 mM trisodium citrate, 30 mM borate with varying pH values was supplemented with or without 128 mM CaCl_2_. p*Ka* values were then obtained with previously described calculations and curve-fittings performed by Prism 9.5 software employing specific-binding-with-Hill-slope function^2^. Error bars represent SEM.

***Cell isolation and cultures***

*Culture of non-excitable cell lines:* HEK293, HeLa and COS-7 cells (ATCC, cat#: CRL-1573, CL-0101 and CRL-1651, respectively) were grown at 37 °C in a humidified atmosphere containing 5 % CO_2_ in DMEM (HyClone, Chicago, IL, USA) with 10 % fetal bovine serum (cat: FBSSA500-S, AusGeneX, Australia), 5 % penicillin and streptomycin (Thermo Scientific, Waltham, MA, USA).

*Primary dissociated hippocampal neuronal cultures:* The cultured of primary hippocampal neurons involves a meticulous process starting with the dissection of hippocampi from both sides were dissected from embryonic day 18 CD®(SD) IGS rats of either gender using dissection tools previously sterilized by autoclaving. The dissected tissues are treated with a 0.25 % trypsin solution (Sigma‒Aldrich) at 37 °C for 15 minutes, followed by gently shaking gentle shaking at 5-minute intervals to ensure thorough digestion. Subsequently, the dissected tissues were washed with Hanks’ balanced salt solution (HBSS) without Ca^2+^ and Mg^2+^ (Thermo Fisher) to remove debris and blood, followed by pipetting with a Pasteur glass tube (15 mm) in DMEM-F12 (with L-glutamine +10% FBS) complete medium (Thermo Fisher) to obtain a cell suspension. The dissociated neurons were plated on 50k cells/18 mm coverslips in 12-well plates, which were coated overnight with 0.01 % poly-L-lysine (Sigma‒Aldrich, Cat# P1274) at 37 °C. On the first day of culture, the neurons were incubated in DMEM-F12 (with L-glutamine +10% FBS) complete medium in a 5 % CO_2_ atmosphere at 37 °C to facilitate initial neuron adhesion and viability in the optimal developmental growth conditions. After 24 hours of plating, half of the plating medium was replaced with fresh neurobasal medium without serum (with 2 % B27, 0.5 mM L-glutaMAX, 37.5 mM NaCl) and once every 4 days thereafter. Neurons are prepared for experimental use following a development period of 18-20 days in culture.

*Neonatal and Adult rat* *cardiomyocytes isolation and culture:*

Neonatal ventricular myocytes were isolated from 1-day-old Sprague-Dawley rats, as described previously^12^. 0.1 % typsin and 0.08 % collagenase II in Hanks solution was used to digestion. Myocytes were plated at 5 × 10^5^ cells / 3.5 cm dish in DMEM (Invitrogen) supplemented with 10 % FBS (Gibco) in the presence of 0.1 mM 5-bromo-2-deoxyuridine (Sigma).

Adult ventricular cardiomyocytes were enzymatically isolated as previously described^13^. Briefly, hearts were rapidly removed from anesthetized adult Sprague-Dawley rat and perfused using Ca^2+^-free Tyrode’s solution (in mM: 120 NaCl, 4 KCl, 1.2 MgCl_2_, 1.5 NaH_2_PO_4_, 20 NaHCO_3_, 10 glucose, 30 taurine, 95 % O_2_ / 5 % CO_2_ saturated) in a Langendorff system. Collagenase Type II (0.7 mg/mL) (Worthington Biochemical) was used to digestion of the heart into single cells. After digestion, the myocardium tissue was cut into small pieces and filtered into Ca^2+^-free Tyrode’s solution with a sieve, then replaced with gradient Ca^2+^ solution, resulting in a final solution with 1 mM Ca^2+^. Freshly isolated cardiomyocytes were plated on laminin-coated (Sigma) culture dishes for 20 min and then the attached cells were maintained in M199 medium (Gibco) at 37 °C under 5 % CO_2_ until usage.

***Gene transfections***

*Gene transfection animal cell lines:* Gene transfection of cells was completed with electroporation (Bio-Rad Gene Pulser Xcell system, Hercules, CA, USA). The electroporation protocol was a voltage step pulse (180 V, 25 ms) using 4 mm cuvettes and 0.4 mL OPTI-MEM medium. Transfected cells were seeded on round coverslips and cultured with DMEM. Imaging experiments were carried out 24 h after transfection.

*Transfection for primary dissociated hippocampal neurons:* Cultured neurons were transfected with 0.5-1 µg plasmids G-CEPIA1er or NEMOer -encoding plasmids at approximately 10-11 DIV using a calcium phosphate transfection kit (Takara, Cat# 631312). Specifically, cultured neurons on coverslips were transferred to a new 12-well plates with pre-warmed DMEM-F12 (with L-glutamine + 25% 1 M HEPES) complete medium. Mix plasmids with 10 % of 2 mM CaCl_2_ in ddH_2_O. Subsequently, the DNA-containing mixed solution was transferred eight times into 2×HBS solution, vortexing for 3 seconds after each transfer. The mixture was then incubated for 15–20 minutes at room temperature and added dropwise to the culture 12-well plates with medium. When an evenly distributed layer of precipitate particles was observed across the neurons (approximately 2 hours later), the DNA-Ca^2+^-phosphate precipitates were dissolved using 1×HBS solution (pH 6.8). The transfected neurons were then transferred back to their original 12-well plates containing neurobasal medium.

*Gene transfection of rat cardiomyocytes:* For adenovirus-mediated gene expression of adult cardiomyocytes. Cells were infected with adenoviruses carrying the NEMOer-f calcium sensor at a multiplicity of infection (m.o.i.) of 30. Confocal imaging was performed 48 h after virus infection. As to neonatal rat cardiomyocytes, 36 to 48 hr after cell culture, 4 μg plasmids was transiently transfected into cells with Lipofectamine 3000 according to the manufacturer’s instructions.

***Fluorescence imaging***

The fluorescence imaging was performed using a ZEISS observer Z1 imaging system controlled by Slidebook 6.0.23 software (Intelligent Imaging Innovations, Inc.)^2^. Cells grown on coverslips were immersed in a physiological salt solution (107 mM NaCl, 7.2 mM KCl, 1.2 mM MgCl_2_, 11.5 mM glucose, 20 mM HEPES-NaOH at pH 7.2), supplemented with 0.1 % BSA. The excitation/emission filters used: NEMOer (500 ± 12 / 542 ±13 nm), G-CEPIA1er/ER-GCaMP6-150 (470 ± 11 /510 ± 14 nm) and TurN/TuNer (For YFP channel, 500 ± 10 / 535 ± 15 nm; for CFP channel, 438 ± 12 /470 ± 12 nm). Fluorescence was collected every two seconds, and the mean fluorescent intensity of the corresponding regions was exported for analysis using Matlab 2023b (The MathWorks, Natick, MA, USA), and the results were plotted using Prism 9.5 software. Error bars denote SEM.

***Confocal imaging***

*Confocal imaging of COS-7 cells.* Recordings were undertaken with a ZEISS LSM880 microscope equipped with 63x oil objective (NA 1.4) and ZEN 2.1 software. For measurement of TuNer-m fluorescence ratio excited by optimal excitation with and without calcium, excitation was set at 458 and 514 nm, with the emission collected at 460-500 nm and 520-600 nm. The acquired images were analyzed using Image J and Matlab 2023b software. To improve image resolution and contrast, the software MicroscopeX FINER(Computational Super-resolution Biotech Co.,Ltd, Guangzhou, China), which uses sparse deconvolution, was utilized, following the methodology reported previously ^14^.

*Confocal line-scan imaging of cardiomyocytes.* The imaging was performed on confocal microscope (LSM-710; Carl Zeis) equipped with an argon laser (488 nm) and 40×, 1.0 N.A. water immersion objective. Myocytes were loaded with 10 μM Rhod2-AM (Invitrogen) under 37 °C for 10 min, and the experiments were performed within 1 hr after loading. In pacing Ca^2+^ transient recording experiments, 15 V, 1 Hz field stimulation to the myocytes. For Ca^2+^ transient, Ca^2+^ spark and SR load measurement, line-scan images were acquired at sampling rates of 3.78 ms/line, the excitation was at 488 and 543 nm and their fluorescence emission were collected at 490 to 550 and 560 nm to 800 nm. The fluorescence-time curves of image were analyzed by python and IDL program, the ΔF/F_0_ ((F-F_0_)/F_0_), time to peak (the time of F_max_) and τ_90_ (decay time of 90% fluorescence change of F_max_ ) were calculated from curve by matlab R2018a program.

*Ca^2+^ imaging and quantitative analysis of fluorescence intensity in neurons*

Neurons of DIV 18-20 were imaged using a W1 spinning disc confocal microscope (Ti2-E, NIKON) with a 100 × oil-immersion objective (1.45 NA, NIKON) in extracellular solution (120 mM NaCl, 3 mM KCl, 10 mM glucose, 10 mM HEPES-NaOH, 2 mM MgCl_2_, and 2 mM CaCl_2_, pH 7.4). Field stimulations were performed in a stimulation chamber (Warner Instruments, RC-49MFSH) with a programmable stimulator (Master-8, AMPI). The neurons were imaged with an exposure time of 200 ms with consecutive and uninterrupted acquisition. For pharmacological experiments, 50 μM CPA (MCE, Cat# HY-N6771) was incubated for 1 hours in neurobasal medium at 37 °C. In the control group, neurons were incubated with DMSO (dimethyl sulfoxide, Sigma) for the corresponding time.

All the quantifications of fluorescent signals in neuronal cultures were performed on the secondary dendrites and somata of transfected neurons. A region of interest (ROI) was chosen based on the imaging criteria, e.g. the quality of fluorescent signals, the background level. ROI of dendrites and somata were manually defined in ImageJ for calculation of mean fluorescent signal *Fs*. A similar region near the dendrites and somata of interest was selected for calculation of fluorescent background *Fb*. The effective signal was calculated as *F* = *Fs* – *Fb*. Since all our quantifications were based on *F* normalized with the averaged fluorescence intensity within the region during the baseline period, the area of defined polygon would have no impact on the normalized *F*. We also simultaneously monitor x-y drift to ensure that the identification of the target area in ImageJ does not deviate from the defined ROI.

***Multimodality Structured Illumination Microscopy (Multi-SIM) imaging***

SIM images were acquired on a Multi-SIM imaging system^15^ (NanoInsights-Tech Co., Ltd.) equipped with a 100 × 1.49 NA oil objective (Nikon CFI SR HP Apo), and a Photometrics Kinetix camera. The SIM images of Cos7 cells expression TuNer-m were taken using low NA GI mode with laser powers of 100 mW (488 nm) and 300 mW (405 nm), and exposure times of 2 ms and 30 ms, respectively. Images were then reconstructed using the SIM Imaging Analyser software (NanoInsights-Tech). During the acquisition of images, cells were in a humidified chamber maintained at 37 ℃ in the presence of 5% CO_2_. The reconstructed images were analyzed using Image J software.

***Animals***

CD®(SD) IGS rats were obtained from Charles River and employed for the cultured primary hippocampal neuron. The rats were maintained group-housed in cages of three each under 12-hour light/dark cycle conditions (lights on at 7 am) in a stable environment (temperature from 22 °C to 24 °C and water and food available ad libitum). Female CD®(SD) IGS rats were sacrificed at 18 days of gestation for hippocampal neuron culture. All animal experiments were approved by the University of Science and Technology of China of the Animal Care and Use Committee requirements (approval no. USTCACUC26120223022).

***Experiment in zebrafish (Danio rerio)***

To generate transient expression larvae for imaging, mating pairs of adult fish of Nacre background were set up to avoid dark pigmentation. Injection mix contained 25 ng/uL of pTol2-β-actin: NEMOer-f and 25 ng/uL of Tol2 mRNA. Approximately 1 nL of the cocktail was injected into fertilized eggs. Adult and larval zebrafish were maintained at 28 degrees on a 14 hour/10 hour light/dark cycle. All experiments were conducted in 10% Hank’s solution containing (in mM): 140 NaCl, 5.4 KCl, 0.25 Na_2_HPO_4_, 0.44 KH_2_PO_4_, 1.3 CaCl_2_, 1.0 MgSO_4_, and 4.2 NaHCO_3_ (pH 7.2). All experimental procedures were approved by Center for Excellence in Brain Science and Intelligence, Chinese Academy of Sciences.

Confocal microscopy was performed using Olympus FV3000 upright microscope with a LUMPlanFL N 40x water immersion objective (N.A. 0.8, W.D. 3.3). Scanning speed was 1 Hz and a single plane was imaged for time-lapse imaging. Frame size was set as 512 x 512 at 16-bit depth and the actual scan size was slightly cropped to allow 1 Hz speed. The 5 dpf or 6 dpf zebrafish larvae were embedded sideways in 2% low-melting agarose (Invitrogen, 16520100) under system water. Zebrafish larvae were neither anesthetized nor paralyzed. Time-lapse images were registered using StackReg to correct mild drifts and fluorescence values of ROI were measured in Fiji. Baseline fluorescence (F_0_) was defined as the top fifteen percentile value and normalized fluorescence changes (ΔF/F_0_) were calculated.

**Supplementary Table 1.** *In cellulo* screening results of NEMOer candidates ^$^

| No | Name^1^ | Mutations in CaM | L2^2^ | L3^3^ | F_0_ (a.f.u.) | Dynamic range | N^4^ | Primer^5^ |
| --- | --- | --- | --- | --- | --- | --- | --- | --- |
|  | **NEMOs** | template | GGSGGGSSS | IYF | -- | -- | -- |  |
| 1 | **G-CEPIA1er** | -- | -- | -- | 4129±303 | 8.5±0.4 | 105/67 |  |
| 2 | **NCaMPer** | E31D-F92W-E104D-D133E | GGSGGGSSS | MYF | 7217±968 | 2.3±0.2 | 16/16 | 1-12 |
| 3 | **NCaMPer1** | E31D-F92W-E104D-D133E | G | MYF | 10374±747 | 32.6±1.7 | 79/79 | 1,12,13,14 |
| 4 | **NCaMPer2** | E31D-F92W-E104D-D133E | GG | MYF | 8655±477 | 30.5±1.7 | 81/81 | 1,12,13,15 |
| 5 | **NCaMPer3** | E31D-F92W-E104D-D133E | GGS | MYF | 7334±446 | 26.3±1.7 | 96/96 | 1,12,13,16 |
| 6 | **NCaMPer4** | E31D-F92W-E104D-D133E-E140D | G | MYF | 11383±1086 | 48.4±1.5 | 152/27 | 1,12,17,18 |
| 7 | **NCaMPer5** | E31D-F92W-E104D-D133E-E140D | GG | MYF | 13208±683 | 63.8±2.5 | 169/38 | 1,12,17,18 |
| 8 | **NCaMPer6** | E31D-F92W-E104D-D133E-E140D | GGS | MYF | 15948±827 | 49.0±1.0 | 195/51 | 1,12,17,18 |
| 9 | **NCaMPer4F** | Q3D-A15E-E31D-M36F-F92W-E104D-D133E- E140D | G | MYF | 1400±106 | 3.2±0.2 | 43/43 | 1,12,19-22 |
| 10 | **NCaMPer5F** | Q3D-A15E-E31D-M36F-F92W-E104D-D133E- E140D | GG | MYF | 1231±118 | 13.7±0.5 | 37/37 | 1,12,19-22 |
| 11 | **NCaMPer6F** | Q3D-A15E-E31D-M36F-F92W-E104D-D133E- E140D | GGS | MYF | 1485±76 | 13.2±0.6 | 48/48 | 1,12,19-22 |
| 12 | **NCaMPer7** | E31D-E67D-F92W-E104D-D133E-E140D | G | MYF | 1302±62 | 3.8±0.2 | 31/31 | 1,12,23,24 |
| 13 | **NCaMPer8** | E31D-E67D-F92W-E104D-D133E-E140D | GG | MYF | 545±58 | 24.4±3.2 | 22/22 | 1,12,23,25 |
| 14 | **NCaMPer9** | E31D-E67D-F92W-E104D-D133E-E140D | GGS | MYF | 972±73 | 24.2±1.1 | 34/34 | 1,12,23,26 |
| 15 | **NCaMPer-L5** | E31D-F92W-E104D-D133E-E140D | GGGGS | MYF | 16558±1265 | 18.5±0.9 | 41/41 | 1,12,13,25 |
| 16 | **NCaMPer-L7** | E31D-F92W-E104D-D133E-E140D | GGGSGGS | MYF | 15380±1047 | 54.4±2.4 | 42/42 | 1,12,13,26 |
| 17 | **NCaMPer-L5F** | Q3D-A15E-E31D-M36F-F92W-E104D-D133E- E140D | GGGGS | MYF | 3362±446 | 21.4±0.8 | 36/36 | 1,12,13,26 |
| 18 | **NCaMPer-L7F** | Q3D-A15E-E31D-M36F-F92W-E104D-D133E- E140D | GGGSGGS | MYF | 314±51 | 35.0±2.2 | 29/29 | 1,12,13,26 |
| 19 | **NCaMPer6-A15E** | A15E-E31D-F92W-E104D-D133E-E140D | GGS | MYF | 2303±159 | 16.3±0.8 | 42/42 | 1,12,27,28 |
| 20 | **NCaMPer3-M/I (NEMOer-m)** | E31D-F92W-E104D-D133E | GGS | IYF | 27665±729 | 141.5±4.5 | 463/122 | 1,12,29.30 |
| 21 | **NCaMPer4-M/I** | E31D-F92W-E104D-D133E-E140D | G | IYF | 2004±84 | 57.5±1.8 | 272/87 | 1,12,29.30 |
| 22 | **NCaMPer5-M/I (NEMOer-n)** | E31D-F92W-E104D-D133E-E140D | GG | IYF | 1515±70 | 177.4±8.1 | 240/83 | 1,12,29.30 |
| 23 | **NCaMPer6-M/I** | E31D-F92W-E104D-D133E-E140D | GGS | IYF | 4920±208 | 107.3±3.9 | 263/86 | 1,12,29.30 |
| 24 | **NCaMPerL5-M/I (NEMOer-s)** | E31D-F92W-E104D-D133E-E140D | GGGGS | IYF | 19371±684 | 138.7±6.6 | 310/111 | 1,12,29.30 |
| 25 | **NCaMPerL7-M/I (NEMOer-b)** | E31D-F92W-E104D-D133E-E140D | GGGSGGS | IYF | 32116±1263 | 67.1±1.7 | 309/104 | 1,12,29.30 |
| 26 | **NCaMPer8-M/I** | E31D-E67D-F92W-E104D-D133E-E140D | GG | IYF | 346±33 | 10.4±0.5 | 71/33 | 1,12,29.30 |
| 27 | **NCaMPer9-M/I** | E31D-E67D-F92W-E104D-D133E-E140D | GGS | IYF | 556±43 | 6.4±0.5 | 90/31 | 1,12,29.30 |
| 28 | **NCaMPer3-M/I-AE** | E14A-A15E-E31D-F92W-E104D-D133E | GGS | IYF | 1452±119 | 28.9±0.8 | 125/50 | 1,12,31,32 |
| 29 | **NCaMPerL5-M/I-AE** | E14A-A15E-E31D-F92W-E104D-D133E-E140D | GGGGS | IYF | 535±33 | 10.7±0.3 | 96/34 | 1,12,31,32 |
| 30 | **NCaMPer2-MI (NEMOer-c)** | E31D-F92W-E104D-D133E | GG | IYF | 25218±733 | 314.5±9.3 | 505/215 | 1,12,33,34 |
| 31 | **NCaMPerL5f-DE-MI (NEMOer-f)** | Q3D-A15E-E31D-M36F-F92W-E104D-D133E | GGGGS | IYF | 10486±340 | 54.7±1.3 | 431/208 | 1,12,29,30,33,34 |
| 32 | **NCaMPerL7f-DE-MI** | Q3D-A15E-E31D-M36F-F92W-E104D-D133E | GGGSGGS | IYF | 12064±433 | 50.8±2.0 | 384/147 | 1,12,29,30,33,34 |
| 33 | **NCaMPerL9-MI** | E31D-F92W-E104D-D133E-E140D | GGSGGGSSS | IYF | 36119±538 | 55.5±2.8 | 239/61 | 1,12,13,35 |
| 34 | **NCaMPerL9F-DE-MI** | Q3D-A15E-E31D-M36F-F92W-E104D-D133E | GGSGGGSSS | IYF | 20445±1002 | 37.0±1.6 | 28/28 | 1,12,13,35 |

Note:

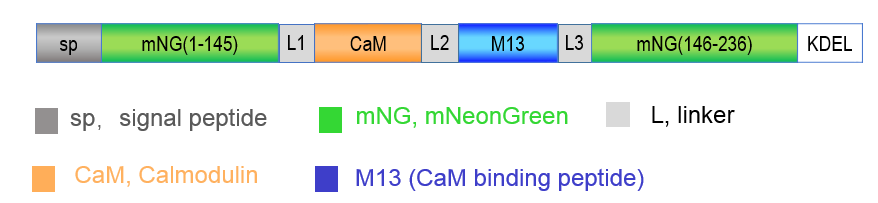
^$^, diagram showing the design of candidate GECIs based on NEMO

--: There is no relevant experiment were performed in this study.

/: NA.

^1^ KOZAK: KOZAK consensus sequence was used.

^2^ L2, Linker between CaM and M13.

^3^ L3, Linker between M13 and mNG.

^4^ N=A/B: A, number of cells used for measurements of the basal fluorescence，B, number of cells used for calculating the dynamic ranges

^5^Primers used, See Supplementary Table 7 for primer sequences corresponding to numbering.

**Supplementary Table 2.** *In vitro* Ca^2+^ titration results and kinetics of NEMOer variants

| Sensor | K_d_ (μM) | Hill_n_ | F_min_  (0 mM Ca^2+^) | F_max_  (128 mM Ca^2+^) | Dynamic range  (ΔF/F_min_) | *k_off_* (s^-1^) |
| --- | --- | --- | --- | --- | --- | --- |
| G-CEPIA1er | 666±44 | 2.0±0.2 | 400±4 | 3456±17 | 6.4±0.1 | 30.57±8.9 |
| NEMOer-m | 922±26 | 1.8±0.1 | 123±0 | 24558±246 | 198.8±1.8 | 17.80±0.3 |
| NEMOer-s | 2599±104 | 1.4±0.1 | 149±2 | 25921±187 | 175.6±1.4 | 11.84±0.2 |
| NEMOer-c | 1203±77 | 1.6±0.1 | 102±1 | 29668±252 | 294.2±2.3 | 19.86±0.3 |
| NEMOer-f | 952±30 | 1.5±0.1 | 93±1 | 11404±70 | 120.5±1.1 | 33.13±0.8 |
| NEMOer-b | 1305±34 | 1.5±0.1 | 210±3 | 30819±166 | 145.4±1.5 | 9.72±0.2 |

**Supplementary Table 3.** *In vitro* biophysical properties of NEMOer sensors

| Sensor | Ca^2+^ | ρ^*1^ | | ε_max_ (mM^-1^*cm^-1^) | | | Φ | | F_1_^*2^ (mM^-1^*cm^-1^) | | p*Ka* |
| --- | --- | --- | --- | --- | --- | --- | --- | --- | --- | --- | --- |
|  |  | Anionic | Neutral | Anionic | | Neutral | Anionic | Neutral | Anionic | Neutral |  |
| G-CEPIA1er | + | 0.13±0.01 | 0.87±0.00 | 71.09±3.39 | 33.21±2.00 | | 0.49 | 0.10 | 4.54±0.37 | 2.89±0.15 | 7.40 |
|  | - | 0.04±0.01 | 0.96±0.01 | 75.49±11.77 | 44.28±1.31 | | 0.24 | 0.13 | 0.62±0.08 | 5.54±0.17 | / |
| NEMOer-c | + | 0.75±0.01 | 0.25±0.02 | 121.50±4.97 | 49.28±1.31 | | 0.60 | 0.16 | 54.85±2.19 | 1.96±0.07 | 5.77 |
|  | - | 0.24±0.01 | 0.76±0.01 | 7.57±1.40 | 69.73±6.14 | | 0.09 | 0.05 | 0.16±0.03 | 2.60±0.24 | / |

Note: ^*1^ ρ is the relative concentration of chromophore; ^*2^ F_1_ is one-photon brightness defined as the product of ρ, ε and Φ.

**Supplementary Table 4.** *In cellulo* screening results of TurN7^$^.

| No. | Name^1^ | Donor | Linker | Acceptor | Basal ratio^2^ | Changes in the donor emission (%) | NCaMP7 Dynamic Range(ΔF/F_min_) | Dynamic range  (ΔR/R_min_) | N^3^ | Primer^7^ |
| --- | --- | --- | --- | --- | --- | --- | --- | --- | --- | --- |
| **1** | **TurN0.1** | mTq2 | - | NCaMP7 | 0.071±0.003 | 41.1±0.4 | 20.9±0.7 | 31.2±1.0 | 33/33 | 36,37,38,39 |
| **2** | **TurN0.2** | mTq2 | G | NCaMP7 | 0.071±0.009 | 33.8±0.6 | 21.5±1.0 | 31.1±1.7 | 28/28 | 36,37,38,40 |
| **3** | **TurN0.3** | mTq2 | GG | NCaMP7 | 0.078±0.004 | 36.4±0.5 | 18.7±0.7 | 27.5±1.1 | 43/43 | 36,37,38,41 |
| **4** | **TurN0.4** | mTq2 | GGS | NCaMP7 | 0.073±0.002 | 32.3±0.5 | 20.1±0.5 | 30.9±0.8 | 40/40 | 36,37,38,42 |
| **5** | **TurN0.5** | mTq2 | GGGS | NCaMP7 | 0.078±0.003 | 32.3±0.3 | 19.4±0.7 | 26.8±1.0 | 40/40 | 36,37,38,43 |
| **6** | **TurN0.6** | mTq2 | GGGGS | NCaMP7 | 0.071±0.003 | 35.6±0.6 | 20.6±0.6 | 31.4±0.9 | 33/33 | 36,37,38,44 |
| **7** | **TurN0.7** | mTq2 | GGSGGS | NCaMP7 | 0.071±0.002 | 34.6±0.3 | 20.2±0.6 | 29.4±0.8 | 33/33 | 36,37,38,45 |
| **8** | **TurN0.8** | mTq2 | GGGSGGS | NCaMP7 | 0.070±0.005 | 35.3±1.0 | 20.6±0.9 | 30.1±1.2 | 24/24 | 36,37,38,46 |
| **9** | **TurN0.9** | mTq2 | GGGGSGGGS | NCaMP7 | 0.081±0.004 | 34.2±1.4 | 17.2±0.5 | 24.2±0.8 | 29/29 | 36,37,38,47 |
| **10** | **TurN1.0** | mTq2 | GGGGSGGGGS | NCaMP7 | 0.081±0.005 | 30.3±1.1 | 18.2±0.6 | 25.4±0.9 | 23/23 | 36,37,38,48 |
| **11** | **TurN1.1** | mTq2 | SGLRSSDPPVAT | NCaMP7 | 0.085±0.007 | 33.1±0.6 | 18.1±1.1 | 24.3±1.5 | 21/21 | 36,37,49,50 |
| **12** | **TurN1.2** | mTq2-ΔC11^4^ | - | ΔN5^5^-NCaMP7 | 0.082±0.002 | 57.2±0.6 | 17.6±0.3 | 40.1±1.3 | 241/105 | 37.51,52,53 |
| **13** | **TurN1.3** | mTq2-ΔC11 | GG | ΔN5-NCaMP7 | 0.072±0.002 | 51.5±0.5 | 19.8±0.3 | 39.8±0.8 | 149/149 | 37.51,52,54 |
| **14** | **TurN1.4 (TurN7)** | sfTq2^ox^-ΔC11 | - | ΔN5-NCaMP7 | 0.083±0.001 | 55.0±0.6 | 16.8±0.6 | 38.4±0.8 | 221/102 | 37.51,52,54 |
| **15** | **CerN^6^0.1** | mCerulean3-ΔC11 | GG | ΔN5-NCaMP7 | 0.101±0.005 | 45.8±0.9 | 17.7±0.5 | 34.6±1.1 | 40/40 | 37.51,52,53 |
| **16** | **CerN0.2** | mCerulean3-ΔC11 | -- | ΔN5-NCaMP7 | 0.097±0.006 | 60.5±0.6 | 18.3±0.8 | 52.9±2.5 | 21/21 | 37.51,52,54 |

Note:

^$^, diagram showing the design of candidate GECI of TurN7.

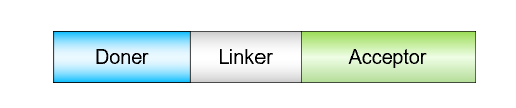

^1^ KOZAK: KOZAK consensus sequence was used.

^2^ Basal ratio: the basal fluorescent intensity ratio of acceptor to donor.

**^3^** N=A/B: A, number of cells used for measurements of the basal ratio，B, number of cells used for calculating the dynamic ranges.

^4^ΔC11, 11 amino acids were truncated from the C-terminus of mTq2-228-238 or sfTq2^ox^-228-238.

^5^ΔN5, 5 amino acids were truncated from the N-terminus of NCaMP7-1-5.

^6^ CerN is composed of mCerulean3-ΔC11 and ΔN5-NCaMP7. However, mCerulean3 is easily quenched than mTq2, hindering further optimization.

^7^ Primers used, See Supplementary Table 7 for primer sequences corresponding to numbering.

**Supplementary Table 5.** *In cellulo* characterization of TurN sensors based on NEMO indicators ^$^.

| Name^1^ | L | Acceptor | Basal ratio | Dynamic range  (ΔR/R_min_) | N^3^ |
| --- | --- | --- | --- | --- | --- |
| TurNc | N/A | ΔN5^2^-NEMOc | 0.0133±0.0003 | 645.3±13.1 | 90/90 |
| TurNm | GGS | ΔN5-NEMOm | 0.0267±0.0006 | 378.4±8.4 | 237/77 |
| TurNf | GGS | ΔN5-NEMOf | 0.0146±0.0002 | 353.1±5.0 | 231/99 |
| TurNb | GGS | ΔN5-NEMOb | 0.0984±0.0031 | 195.0±5.6 | 290/86 |

Note:

^$^, diagram showing the design of candidate GECI of TurN series.

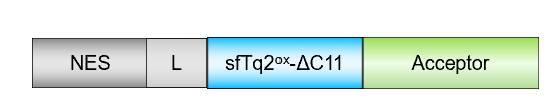

^1^ KOZAK: KOZAK consensus sequence was used.

^2^ΔN5, 5 amino acids were truncated from the N-terminus of mNeonGreen 1-5 in NEMO sensors.

**^3^** N=A/B: A, number of cells used for measurements of the basal ratio，B, number of cells used for calculating the dynamic ranges.

**Supplementary Table 6.** *In cellulo* characterization of TuNer sensors based on NEMOer indicators ^$^.

| Name^1^ | Acceptor | Basal ratio | Dynamic range  (ΔR/R_min_) | K_d_ | Hill_n_ | N |
| --- | --- | --- | --- | --- | --- | --- |
| TuNer-m | ΔN5^2^-NEMOer-m | 1.44±0.05 | 229.8±6.8 | 958.67±32.96 | 1.28±0.05 | 108 |
| TuNer-c | ΔN5-NEMOer-c | 1.60±0.04 | 385.0±4.8 | 1369.96±43.81 | 1.29±0.04 | 98 |
| TurNer-f | ΔN5-NEMOer-f | 0.56±0.02 | 54.3±0.6 | 1198.28±43.81 | 1.16±0.04 | 110 |
| TuNer-s | ΔN5-NEMOer-s | 0.48±0.01 | 222.5±2.7 | 5473.71±43.81 | 0.99±0.02 | 98 |

Note:

^$^, diagram showing the design of candidate GECI of TuNer series.

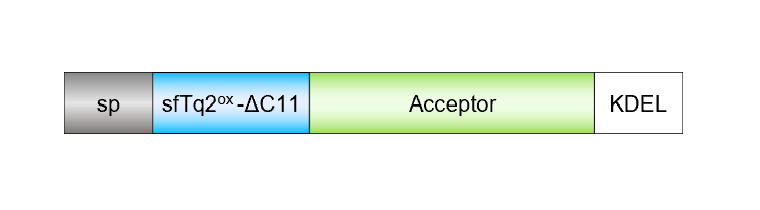

^1^ KOZAK: KOZAK consensus sequence was used.

^2^ΔN5, 5 amino acids were truncated from the N-terminus of mNeonGreen 1-5 in NEMOer sensors.

**Supplementary Table 7.** Sequences of primers used to generate GECI constructs listed in Supplementary Table 1&4.

| No. | Sequence (5’ to 3’) |
| --- | --- |
| 1 | ACCGAGCTCGGATCCGCCACCATGGGATGGAGCTGTATCA |
| 2 | TCGCCCTTGCTCACCATGGATCCCTGCAGTTGGAC |
| 3 | GTCCAACTGCAGGGATCCATGGTGAGCAAGGGCGA |
| 4 | TTACCACAAAAGATCTGGGCACAGTG |
| 5 | GTGCCCAGATCTTTTGTGGTAATGG |
| 6 | ATCAGCGGCGCCAATGTATCCATTGCCGTCCTTGTCCCACACGC |
| 7 | GATACATTGGCGCCGCTGATCTGAGACACGTGAT |
| 8 | ACGGCGAAGGCCAGGTG |
| 9 | CACCTGGCCTTCGCCGTCAAT |
| 10 | GGCAGACTTTCTAGCATGTACTTCGCCGACTGGT |
| 11 | ACCAGTCGGCGAAGTACATGCTAGAAAGTCTGCC |
| 12 | ATATCTGCAGAATTCTTACAGCTCGTCCTTCTTGTACAGCTCGTCCAT |
| 13 | CTTGGCGGTCATCATCTG |
| 14 | CAGATGATGACCGCCAAGGGCAGAAGAAAATGGAAT |
| 15 | CAGATGATGACCGCCAAGGGCGGCAGAAGAAAATGGAAT |
| 16 | CAGATGATGACCGCCAAGGGCGGCAGCAGAAGAAAATGGAAT |
| 17 | ATTATGAGGATTTTGTGCAGATGA |
| 18 | TCATCTGCACAAAATCCTCATAAT |
| 19 | AAGCCCACGACGATCTGACAGAAGAACAGATCGCCGAATTCAAGGAGGAGTTTAGCCTG |
| 20 | CAGGCTAAACTCCTCCTTGAATTCGGCGATCTGTTCTTCTGTCAGATCGTCGTGGGCTT |
| 21 | TGGGCACAGTGTTCAGAAGCCT |
| 22 | AGGCTTCTGAACACTGTGCCCA |
| 23 | GAAAGTCAGGAAAATCCA |
| 24 | ACTGGATTTTCCTGACTTTCTGGC |
| 25 | CAGATGATGACCGCCAAGGGCGGCGGAGGCAGCAGAAGAAAATGGAAT |
| 26 | CAGATGATGACCGCCAAGGGCGGCGGCAGCGGAGGAAGCAGAAGAAAATGGAAT |
| 27 | TCAAGGAGGAGTTTAGCCTG |
| 28 | CAGGCTAAACAGCTCCTTGA |
| 29 | TTTCTAGCATCTACTTCGCCGA |
| 30 | TCGGCGAAGTAGATGCTAGAAAGTC |
| 31 | AATTCAAGGCCGAGTTTAGCCTGTTCGACA |
| 32 | ACAGGCTAAACTCGGCCTTGAATTCGGC |
| 33 | AATTATGAGGAGTTTGTGCAGA |
| 34 | TCTGCACAAACTCCTCATAATT |
| 35 | GGCGGCAGCGGAGGAGGAAGCAGCAGCAGAAGAAAATGGAATAAGGC |
| 36 | ACCGAGCTCGGATCCGCCACCATGGTGAGCAAGGGCGA |
| 37 | CTGGATATCTGCAGAATTCTTACTTGTACAGCTCGTCCAT |
| 38 | CTTGTAAAGCTCGTCCAT |
| 39 | ATGGACGAGCTTTACAAGATGGTGAGCAAGGGCGA |
| 40 | ATGGACGAGCTTTACAAGGGCATGGTGAGCAAGGGCGA |
| 41 | ATGGACGAGCTTTACAAGGGCGGCATGGTGAGCAAGGGCGA |
| 42 | ATGGACGAGCTTTACAAGGGCGGCAGCATGGTGAGCAAGGGCGA |
| 43 | ATGGACGAGCTTTACAAGGGCGGCGGCAGCATGGTGAGCAAGGGCGA |
| 44 | ATGGACGAGCTTTACAAGGGCGGCGGCGGCAGCATGGTGAGCAAGGGCGA |
| 45 | ATGGACGAGCTTTACAAGGGCGGCAGCGGAGGAAGCATGGTGAGCAAGGGCGA |
| 46 | ATGGACGAGCTTTACAAGGGCGGCGGCAGCGGAGGAAGCATGGTGAGCAAGGGCGA |
| 47 | GACGAGCTTTACAAGGGCGGCGGCGGCAGCGGAGGAGGCAGCATGGTGAGCAAGGGCGA |
| 48 | GGCGGCGGCGGCAGCGGAGGAGGCGGAAGCATGGTGAGCAAGGGCGA |
| 49 | GACGGGAGGATCGCTACTTCGAAGACCACTCTTGTAAAGCTCGTCCAT |
| 50 | AGCGATCCTCCCGTCGCTACTATGGTGAGCAAGGGCGA |
| 51 | GGTACCGAGCTCGGATCCGCCACCATGCATCATCATCATCATCATGTGAGCAAGGGCGA |
| 52 | GGCGGCGGTCACGAACTC |
| 53 | GAGTTCGTGACCGCCGCCGAGGAGGAGAACATGG |
| 54 | GAGTTCGTGACCGCCGCCGGCGGCGAGGAGGAGAACATGG |

**Extended Data Figure Legends**

**Extended Data Figure 1. *In vitro* characterization of NEMOer sensors.**

(A) *In vitro* dose–response curves of GECIs. Left, typical traces; right, statistics. (n=3 independent replicates). Data shown as mean ± s.e.m.

**(**B) Kinetics of GECIs’ off-responses. Left, representative normalized fluorescence traces; right, statistics. (n=3 independent replicates). Data shown as mean ± s.e.m.

(C-G) Spectral properties of NEMOer-c and G-CEPIA1er. (A-B) At pH7.2, typical traces of excitation spectrum, emission spectrum (C), and absorption spectrum (D). (E-G) pH-dependence of absorption spectrum at the apo state (E) and Ca^2+^ saturated conditions (F). (G) Statistics for data shown in panels E&F. n=3 independent biological replicates.

**Extended Data Figure 2. Photobleaching properties of NEMOer and G-CEPIA1er in HEK293 cells.**

(A) Widefield photobleaching curves. GFP excitation light (470±11 nm) was used. Top, two panels on the left, GCEPIA1er and NEMOer-m signals; right, NEMOer-m fluorescence intensity excited under different light intensities.

(B) Statistics showing the relative reduction in fluorescence at the end of 200 sec illumination with medium (left) or strong (right) light. Left (G-CEPIA1er, n=27 cells; NEMOer-s, n=18 cells; NEMOer-f, n=40 cells; NEMOer-c, n=30 cells; NEMOer-b, n=27 cells; NEMOer-m, n=27 cells); right (G-CEPIA1er, n=31 cells; NEMOer-s, n=19 cells; NEMOer-f, n=40 cells; NEMOer-c, n=30 cells; NEMOer-b, n=27 cells; NEMOer-m, n=27 cells). Three independent biological replicates.

**Extended Data Figure 3. Performance of miGer and TuNer indicators in HEK293 cells.**

**(**A) Diagram illustration of the design of TuNer indicator. The CFP variant, sfTq2^ox^, attached at the N-terminus of NEMOer-ΔN5 serves both as a reference fluorescence protein and FRET donor. Due to alterations in FRET efficacy, its fluorescence changes reciprocally with that of NEMOer.

(B) Typical traces showing the TuNer-m responses indicated by fluorescence of NEMOer-m (F_NEMOer-m_, bottom left), sfTq2^ox^ (F_sfTq2ox_, top left), along with the corresponding fluorescence ratio (R), or F_NEMOer-m_ / F_sfTq2ox_.

(C) GECI responses induced by 10 μM CCh. Top, typical traces; bottom, statistics (miGer, n=109 cells; TuNer-s, n=104 cells; TuNer-f, n=115 cells; TuNer-c, n=98 cells; TuNer-m, n=104 cells) (*****P* < 0.0001, One-way ANOVA).

(D) Statistics of basal ER Ca^2+^ levels of HeLa (top) or HEK293 (bottom) cells measured by miGer or TuNer indicators. Top (miGer, n=97 cells; TuNer-s, n=98 cells; TuNer-f, n=110 cells; TuNer-c, n=98 cells; TuNer-m, n=108 cells); bottom (miGer, n=62 cells; TuNer-s, n=130 cells; TuNer-f, n=82 cells; TuNer-c, n=91 cells; TuNer-m, n=141 cells).

Data in (C) and (D) were shown as mean ± s.e.m.

**Extended Data Figure 4.** **Ratiometric signals of TuNer-m transiently expressed in COS-7 cells.**

(A) Typical basal fluorescence or ratiometric image. Regions indicated by white arrows show nuclear ER, while regions indicated by yellow arrows show ER adjacent to the nucleus. Scale bar, 10 μm.

(B) The enlarged portion of the cell from (A). Regions indicated by white arrows show higher NEMOer-m signal but similar TuNer-m ratio, as compared with their corresponding adjacent areas. Scale bar, 10 μm.

(C) Statistics the ratio of TuNer-m in the nuclear ER regions and ER adjacent to the nucleus regions. (****** *P* < 0.0001, paired Student’s t-test, two-tailed.)

(D) Statistics of subcellular regions with high or low TuNer-m ratio. (****** *P* < 0.0001, paired Student’s t-test, two-tailed.)

(E) Typical subcellular TuNer-m responses in COS-7 cells stimulated with 5 μM ATP. (n = 3 independent biological replicates, with at least 12 cells per repeat).

**Extended Data Figure 5. Performance of NEMOer-f in adult rat cardiomycytes.**

Cells transfected with NEMOer-f were loaded with Rhod-2, and dual-color line-scan imaging were performed to simultaneously monitor cytosolic and SR Ca^2+^ changes. Three independent biological replicates.

(A) Ca^2+^ responses trigged by a train of 1Hz electric field stimulation. Top, typical line-scan images; bottom, corresponding mean time course plots.

(B) A replot of Fig. 4E to better visualize the differences of responses between CT and NE-treated cells.

(C) Statistics of responses in (B) and Fig. 4B. Top, Rhod-2 signals; bottom, NEMOer-f signals (* *P*=0.041, ** *P*=0.068, ****P*=0.0001, *****P*<0.0001, top ns *P=*0.5602, bottom ns *P=*0.9838, unpaired Welch's t test, two-tailed.) (For CT, 11 cells from 3 adult rats; for NE, 20 cells from 3 adult rats.).

**Movie legends**

**Supplementary Video 1. Typical** **ATP-induced ER Ca^2+^ oscillations shown by TuNer-m in a COS-7 cell.** TuNer-m responses to 5 μM ATP stimulation were monitored using confocal microscopy. Scale bar, 10 μm. Regions indicated by purple arrows show higher fluorescence signals but lower TuNer-m ratio compared to their corresponding adjacent areas. Those marked by yellow arrows show higher ratios but lower NEMOer fluorescence compared to their adjacent areas.

**Supplementary Video 2. Zoom-in video 1 of the cell shown in Video 1.** Dots marked by white arrows represent microdomains where NEMOer-m fluorescence either remained consistently higher or transiently increased compared to their neighboring regions, while the corresponding TuNer-m ratios remained similar to adjacent areas. Scale bar, 3 μm.

**Supplementary Video 3. Zoom-in video 2 of the cell shown in Video 1.** TuNer signals during the formation of an ER tubule was indicated by white arrows. “Moving dark dots” of TuNer-m ratio, or transient lowering of ER Ca^2+^ levels in microdomains within ER tubules are indicated by yellow arrows. Scale bar, 3 μm.

**Supplementary Video 4. Typical *in vivo* spontaneous SR Ca^2+^ releasing events reported by NEMOer-f in muscle cells of zebrafish.**
